# Structure and Dynamics of HIV-1 Env Trimers on Native Virions Engaged in Living T Cells

**DOI:** 10.1101/2020.08.16.252742

**Authors:** Irene Carlon-Andres, Tomas Malinauskas, Sergi Padilla-Parra

## Abstract

The HIV-1 envelope glycoprotein (Env) mediates viral entry into the host cell. Although the highly dynamic nature of Env intramolecular conformations has been shown with single molecule spectroscopy in vitro, the bona fide Env intra- and intermolecular mechanics when engaged in live T cells remains unknown. We used both, two photon fast fluorescence lifetime imaging detection of single-molecule Förster Resonance Energy Transfer and single molecule photobleaching to reveal transitions between intramolecular and intermolecular conformations that mediate Env clustering. Furthermore, we show that three broad neutralizing anti-Env antibodies directed to different epitopes destabilise Env intramolecular dynamics and their clusters when engaged to living T cells. Importantly, our results show that Env clustering is a common conformation across different HIV-1 Env strains, which depends on efficient virus maturation, and that is disrupted upon binding of Env to CD4 or to neutralizing antibodies. Thus, this study illuminates how different intramolecular conformations and clusters of Env mediate HIV-1 Env–T cell interactions in real time and therefore might control immune evasion.

## Introduction

HIV-1 entry into the cell requires fusion of the viral lipidic envelope with the host plasma membrane. This process is mediated by the HIV-1 Env glycoprotein, consisting of a homotrimer of gp120-gp41 heterodimers ^1^, which is found in low density per virion (7–14 spikes per particle) ^2–4^. The interaction between HIV-1 Env and host receptor CD4 triggers a series of conformational changes allowing the co-receptor, either CCR5 or CXCR4, to bind to the prefusion complex ^5,6^. We have previously determined the time-resolved stoichiometry of the prefusion complex in live cells ^5^. This prefusion complex assembled via a common three-step mechanism in which one or two Env protomers were engaged in the prefusion complex. The current hypothesis for Env intramolecular dynamics HIV-1 Env would undergo three states during this process: first, Env adopts a closed conformation (named State 1) right before CD4 asymmetric interaction; second, after CD4 engagement, Env adopts an intermediate state (State 2) followed by a last open conformation for the coreceptor engagement (State 3) that exposes otherwise hidden, more conserved epitopes, increasing susceptibility for antibody recognition ^7–9^.

The intramolecular structure and dynamics of the HIV-1 Env has been extensively studied during the past few years ^8–11^. Munro et al., ^9^ pioneered *in vitro* intramolecular structural and dynamic studies of HIV Env trimers in native virions utilizing single molecule Förster Resonance Energy Transfer (FRET) and described three different intramolecular conformational states of Env. It is currently unclear how these three different states can be reconciled with Env diffusion ^3^ and intermolecular dynamics ^12^ during cluster formation and dissociation in mature HIV-1 viruses and its relation with the prefusion reaction on the surface of the host. Moreover, it is still not clear whether these three intramolecular states described *in vitro* recapitulate the bona fide dynamics of HIV-1 Env when engaged in live T cells, in the presence or absence of broadly neutralizing antibodies.

In this study, we were able to detect intramolecular conformational states of Env, also described before ^9^, and to determine how they relate to intermolecular Env cluster conformations during the first steps of the HIV-1 prefusion reaction in living T cells. We also describe the role of Env clusters when exposed to inhibiting concentrations of broadly neutralizing antibodies.

## Results

### Structural Characterization of HIV-1 virions by two photon FRET-FLIM

Aiming to ascertain both intramolecular and intermolecular Env dynamics with a multiparameter FRET and fluorescence lifetime microscopy (FLIM) approach, we produced HIV-1 virions labelled with a variant of super-folding GFP (GFP_OPT_) in the V4 loop of gp120 HXB2 Env glycoprotein (Fig. 1A-C) ^13^. Importantly, it was previously shown that labelling the V4 loop of gp120 with GFP_OPT_ does not significantly interfere with HIV-1 fusogenic activity ^14^. HIV-1 virions pseudo-typed with HXB2 V4-GFP_OPT_ Env were exposed to monoclonal nanobodies against GFP, in turn, labelled with Atto 488 (NbA488) and Atto 594 (NbA594), that constitute the donor and acceptor dipoles of the FRET pair, respectively (Fig. 1C). This particular labelling strategy allows FRET to occur between donor and acceptor dipoles located in a single Env molecule (intramolecular interaction) and between adjacent Env molecules (intermolecular interaction), when fluorophores are in close enough proximity and in a proper orientation. To be certain of only considering bona fide HIV-1 virions and being able to determine their maturation state for subsequent FRET-FLIM analysis, pseudo-virions were produced harbouring the Gag polyprotein precursor fused to GFP (Fig. 1A-B). Virion labelling efficiency was determined by exposing HIV-1_Gag-GFP HXB2 V4-GFPopt_ virions to NbA594 and quantifying the percentage of double positive (GFP+ NbA594+) particles, which was 32.7% of the total GFP+ particles (Fig. S1A). We also assessed the maturation efficiency of the viral sample by tracking the release of the internal envelope GFP after exposure to 0.01% concentration of saponin. Only mature HIV-1 particles that undergo proteolytic processing of Gag are able to release the GFP content after permeabilization of the viral membrane. We observed that 40% of virions were able to release the internal GFP against 0%, when virions were produced in presence of the HIV-1 protease inhibitor Saquinavir (SQV) (Fig. S1B). Double positive (GFP+ NbA594+) mature virions showed a drop in GFP fluorescence upon saponin treatment and a stable Atto594 fluorescence signal overtime (Fig. S1C, top panel), showing both, a low contribution of photons from the V4-GFP_OPT_ Env compared to Gag-GFP, and a high stability of the Atto594 fluorophore, making this labelling suitable for our FRET-FLIM experiments. In contrast, immature virions (+SQV) did not show a drop in GFP fluorescence overtime but an increase in Atto594 fluorescence intensity, suggesting an increased accessibility of the NbA594 to the unprocessed Gag-GFP after viral membrane permeabilization (Fig. S1C, bottom panel).

**Figure 1.**
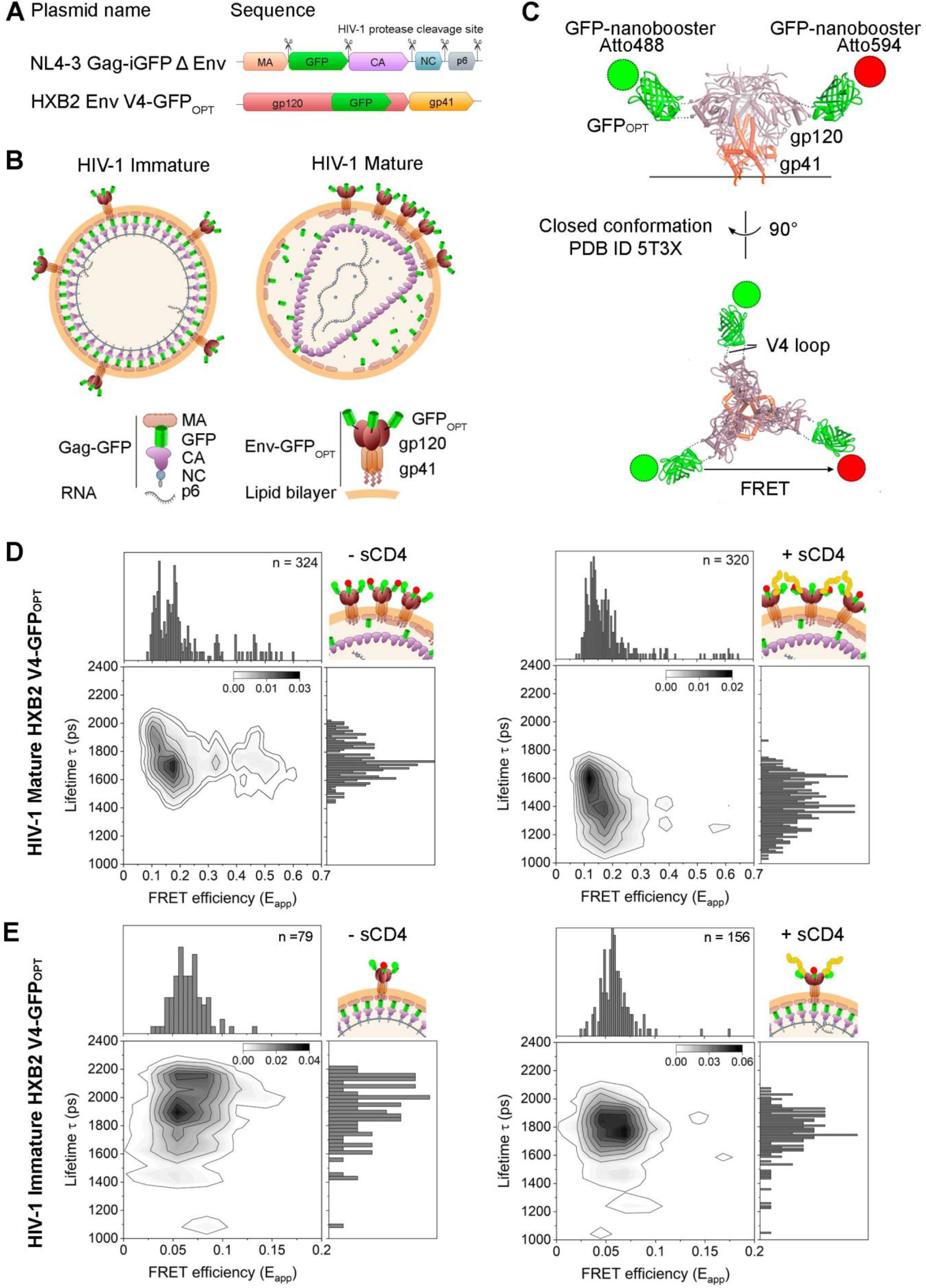
Two Photon FRET-FLIM detects HIV-1 Env conformations. (A) Diagram representing the DNA sequence of the plasmids used to produce HXB2 pseudotyped virions. MA: matrix. GFP: green fluorescent protein. CA: capsid. NC: nucleocapsid. Protease cleavage sites are indicated by scissors symbol. (B) Schematic representation of HIV-1 immature (left) and mature (right) particles used in this assay. Immature particles possess the unprocessed Gag polyprotein fused to GFP. The viral protease cleaves Gag to mediate assembly of the mature HIV-1 virion. Few copies of HXB2 V4-GFP_OPT_ embedded in the viral membrane allow analysis of Env conformations by FRET-FLIM and single-molecule approaches. (C) The HIV-1 Env structure representing closed Env (PBD ID 5T3X) conformation was modified to illustrate the labeling of the V4 loop with GFP_OPT_ and NbA488 and NbA594. (D-E) Two-dimensional (2D) kernel probability graphs showing FRET (FRET efficiency, E_app_) vs FLIM (Lifetime, in ps) data. High density regions are depicted in darker grey color. (D) Plots corresponding to HIV-1 mature HXB2 V4-GFP_OPT_ labelled with NbA488 and NbA594 nanobodies incubated without (left) or with (right) sCD4. (E) Plots corresponding to HIV-1 immature HXB2 V4-GFP_OPT_ labelled with NbA488 and NbA594 nanobodies incubated without (left) or with (right) 10 μg/mL concentration of soluble CD4 (sCD4_D1-D4_).

HIV-1_Gag-GFP HXB2 V4-GFPopt_ virions in presence or absence of Nbs were imaged using two photon rapid FLIM ^15,16^ and both lifetime and apparent FRET efficiency were calculated and plotted into multi-parameter two-dimensional graphs ^17^. The multidimensional analysis of correlated changes of FRET and FLIM allows to efficiently detect heterogeneities in a population even in low-photon conditions, where few labelled proteins are available ^18^. Hence, this technique is able to separate the spectrally overlapped signals of GFP and Atto 488 based on their different intrinsic lifetimes (1200±100 ps and 1800±50 ps, respectively) (Fig. S2A). Clearly, the addition of NbA488 induced a lifetime change in a large population of HIV-1 virions, independently of their maturation state (~80% of the HIV-1 virions presented lifetime values of ~1800 ps) (Fig. S2A-B). Therefore, we could efficiently discriminate HIV-1 particles labelled with NbA488 from NbA488-negative particles. In addition, the FRET efficiency profile of HIV-1 particles labelled only with the donor, Nb488, allowed us to define a no-FRET threshold for mature HIV-1_Gag-GFP HXB2 V4-GFPopt_ particles (E_app_ < 0.1; Fig. S2A, right panel) and for immature viral particles (Eapp < 0.06; Fig. S2B, right panel).

To define the heterogenous Env conformational landscape by FRET-FLIM, we exposed HIV-1_Gag-GFP HXB2 V4-GFPopt_ particles to both, NbA488 and NbA594. If FRET would occur between Env V4 labels (NbA488 and NbA594), the fluorescence lifetime of NbA488 in the presence of NbA594 would be shortened or quenched and the apparent FRET efficiency would be increased ^19,20^. As expected, addition of both, donor and acceptor fluorophores (NbA488 and NbA594, respectively) induced a shift towards positive E_app_ values (E_app_ > 0.1), which correlated with decreasing lifetime values (Fig. 1D, left panel). Here, we could detect three main FRET regimes: i) low or no FRET (E_app_ < 0.12) and higher lifetimes (~1900 ps) ii) a more dense population showing intermediate FRET efficiency (0.12 < E_app_ < 0.23) and moderately decreased lifetimes (~1750 ps), and iii) high apparent FRET efficiency (E_app_ > 0.23) and decreased lifetimes (~ 1700 ps).

Seeking to relate the observed FRET-FLIM profile with intramolecular conformations of HIV-1 Env in our functional virions, we exposed mature HIV-1_Gag-GFP HXB2 V4-GFPopt_ particles, to saturating concentrations (10 μg/mL) of soluble CD4 (sCD4_D1-D4_) (Fig. 1D, right panel). We could readily stabilize a low or no-FRET situation (E_app_ < 0.12) that we could assign to an Env open conformation (Fig. 1D, right panel) as seen by others ^21^, whereas the intermediate and the hight FRET regime populations were clearly reduced. This result suggests that intermediate (0.12 < E_app_ < 0.23) and high FRET regimes (E_app_ > 0.23) could relate to an intramolecular closed Env conformation or intermolecular Env interactions, in which the conditions for FRET to occur would be more favourable.

It has been previously shown that in immature HIV-1 viruses, Env diffuses twice as slow (D = 0.001 μm^2^/sec) as compared to mature HIV-1 particles (D = 0.002 μm^2^/sec) ^3^. Moreover, in immature HIV-1 virions, Env is unable to form clusters ^12^. Based on these observations and to accurately define a FRET threshold for intramolecular interactions, we produced immature HIV-1 virions ^22^ and co-labelled them with nanobodies NbA488 and NbA594. In this case, only two FRET regimes could be determined: i) low or no-FRET (E_app_ < 0.7) and ii) moderate FRET efficiency (0.7 < E_app_ < 0.17) (Fig. 1E, left panel) suggesting that Env in immature viruses adopts at least two conformational states. Addition of saturating concentrations of sCD4_D1-D4_ to immature virions stabilized the low or no-FRET open conformation (E_app_ < 0.7) (Fig. 1E, right panel), as observed in mature virions, showing that immature particles, although impaired for fusion ^12,23,24^, conserve intramolecular dynamics.

When comparing the HIV Env conformations in unbound mature and immature HIV-1_Gag-GFP HXB2 V4-GFPopt_ particles (Fig. 1D and E, left panels), the most prevalent conformation for both was the one showing intermediate FRET efficiencies. It has been previously reported that unligated mature Env preferentially adopts a closed conformation ^9,25^. Therefore, we hypothesized that these moderate FRET values could represent a closed ground-state conformation, as this conformation would reduce the distance between V4 loops within the Env trimer and thus, labels could be close enough to give intramolecular FRET. In turn, both mature and immature virions in presence of sCD4_D1-D4_ (Fig. 1D and E, right panels) showed a predominant low or no-FRET efficiency population, that we could attribute to open Env conformation. Interestingly, high FRET regimes were only observed in mature virions (E_app_ > 0.23), preferentially in the absence of sCD4_D1-D4_ ligand (Fig 1D, left panel). Given that the Env cluster distribution depends on the maturation state of virions ^12^, we attributed hight FRET regimes to intermolecular Env interactions.

To confirm that hight FRET regimes correspond to Env intermolecular interactions, HIV-1 pseudoparticles were produced incorporating the GFP_OPT_ in the V1 loop of gp120 Env glycoprotein instead of the V4 loop (HIV-1_Gag-GFP HXB2 V1-GFPopt_). Note, that this specific labelling strategy was also previously tested for fusogenic Env functionalities ^14^. This labelling approach that positions the donor and acceptor fluorophores proximal to the apex of Env when adopting a closed conformation ^26^, is expected to increase the distance between different Env trimers and therefore drastically reduce or eliminate the intermolecular FRET, if any, between Envs (Fig. S2). Viruses with Env labelled with GFP_OPT_ V1 were exposed to nanobodies coupled to the donor alone (NbA488) or to both, donor and acceptor dipoles (NbA488 and NbA594, respectively). We observed a slight increase in the FRET efficiency in presence of the FRET pair compared to the donor alone condition in both, mature (Fig. S2C) and immature particles (Fig. S2D). However, FRET efficiency was not higher than 0.23, as observed in mature HIV-1_Gag-GFP HXB2 V4-GFPopt_ virions, showing that labelling of the V4 loop in Env is critical to observe Env intermolecular interactions occurring in mature HIV-1 virions.

Therefore, these experiments combining FRET-FLIM to study mature and immature HIV-1 particles, allowed us to discriminate intramolecular (open: E_app_ < 0.12; closed: 0.12 < E_app_ < 0.23) from intermolecular (E_app_ > 0.23) interactions. Of note, while different intramolecular conformations were observed in both, mature and immature particles, intermolecular interactions were only seen in mature HIV-1 virions, suggesting that HIV-1 maturation affects Env clustering but not intramolecular Env conformations. These data also show that sCD4 not only stabilized the open Env conformation, but also disrupted Env clusters as the high FRET regime detected in mature HIV virions was drastically reduced.

### Intermolecular Env conformations are maturation-dependent and modulated by sCD4 and neutralizing antibodies

We further investigated HIV-1 Env cluster formation by single-molecule photobleaching (SMPB) ^27^. This approach relies on the observation of photobleaching dynamics of fluorophores upon continuous illumination. Direct counting of intensity drops from the photobleaching traces reveals the number of fluorescent molecules within the focal volume. In our system, we assumed that constant number of Env glycoproteins are incorporated into HIV-1 virions, irrespectively of the maturation state of viral particles, as previously observed ^12,23,24^. However, we expected to detect relative changes in the number of photobleaching events and/or the overall photobleaching kinetics of mature vs immature viral particles, as a consequence of different Env distribution within the viral membrane and higher probability of synchronous photobleaching of Env molecules within clusters ^27^. In order to be able to spectrally discriminate the Env glycoprotein from the Gag-GFP labelled capsid, mature or immature HIV-1 virions were incubated with NbA594 (Fig. 2A). Photobleaching of the Atto 594 fluorophore was induced by applying a continuous 594 nm laser to the sample and imaged during 350 s. The first 200 s after laser excitation induced the overall intensity decay to drop toward background levels close to zero, suggesting that the bleaching of most of the fluorophores in the sample had occurred (Fig. 2B). Single-drop steps from individual double labelled HIV virions were manually counted from the intensity traces, in both, mature and immature particles, in absence (Fig. 2B) or presence of sCD4_D1-D4_ (Fig. S3A). To exclude any bias in single-drop step quantification, we calculated the mean histogram of all traces per condition and fitted the data to different multi peak Gaussian models; choosing the ones with the reduced Chi-square value closest to 1 (Fig. 2C). We observed a good correlation between the number of resulting Gaussians and the mean of the single-drop step quantification for each condition, which resulted in 5.1±1.1 photobleaching events in case of mature viruses and a statistically significant decrease (3.1±1.1 photobleaching events) for immature viruses (Fig. 2D). Addition of sCD4_D1-D4_ to mature virions caused a slight decrease in the number of photobleaching events observed (4.3±1.1), albeit not statistically significant, and no additive effect in case of immature virions was observed (3.5±1.2).

**Figure 2.**
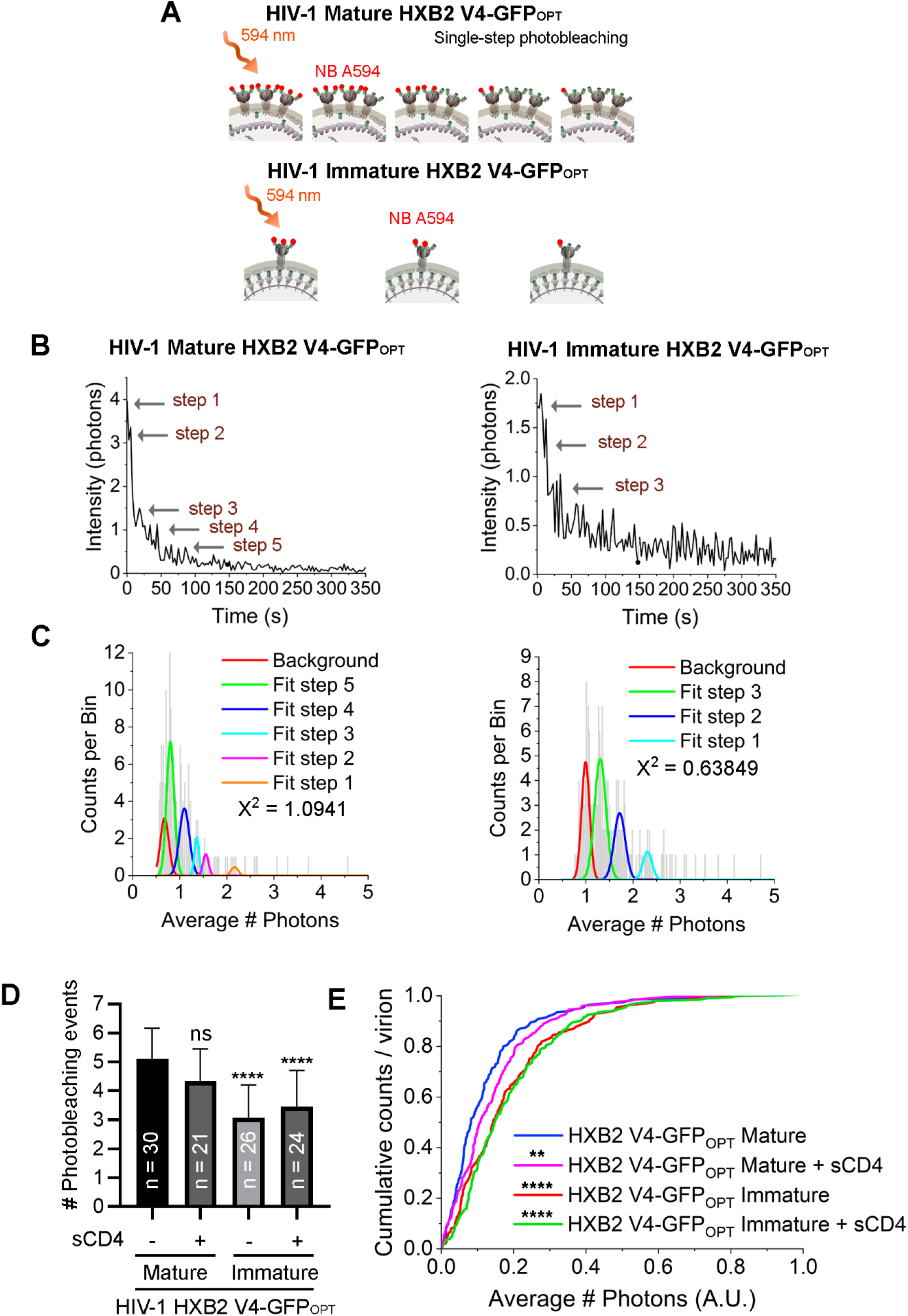
Single-molecule step photobleaching shows differential distribution of HIV-1 Env in mature vs immature HXB2 V4-GFP_OPT_ pseudotyped virions. (A) Schematic representation of the single-molecule step photobleaching approach. Photobleaching of the Atto 594 fluorophore labelling HXB2 V4-GFP_OPT_ was induced by continuous laser 594 nm excitation in mature and immature HIV-1 particles. In case of mature HIV-1 virions, photobleaching of labelled Env is more frequent and faster, suggesting a cluster-like distribution within the viral membrane, as opposed to immature particles, in which photobleaching is less frequent and slower, suggesting a more homogeneous distribution of Env. (B) Representative intensity traces for HIV-1 mature (left) and immature (right) HXB2 V4-GFP_OPT_ particles. Arrows point to single photobleaching steps detected. (C) Population histograms calculated from intensity traces were fitted into a multi-Gaussian distribution model to estimate the number of photobleaching steps for mature (left chart) and immature (right chart) viral particles. Χ^2^ values report the goodness of the fit. (D) Bar graph showing the mean and SD of the number of photobleaching events quantified from intensity traces. N of intensity traces is indicated on each bar per condition. Statistical significance was calculated using a one-way ANOVA and Sidak pos-hoc test comparing each condition to the mature HIV-1 HXB2 V4-GFP_OPT_ condition in absence of sCD4. **** < 0.0001; ns = non-statistically significant (E) Cumulative distribution calculated from the mean intensity trace histograms. Statistical significance was calculated using a Kolmogorov-smirnov test comparing each condition to the mature HIV-1 HXB2 V4-GFP_OPT_ condition in absence of sCD4. ** < 0.01; **** < 0.0001.

To assess whether the photobleaching kinetics of the Atto 594 fluorophore were affected by differences in HIV-1 Env conformation, we compared the cumulative distribution of the intensity traces in mature vs immature virions, in presence or absence of the soluble HIV-1 Env ligand sCD4_D1-D4_. We observed that kinetics of photobleaching are faster in mature HIV-1_Gag-GFP HXB2 V4-GFPopt_ virions compared to immature virions, in a statistically significant manner. Interestingly, binding of sCD4_D1-D4_ to HXB2 V4-GFP_OPT_ Env also induced a delay in photobleaching of Atto 594 in mature viruses, causing no additive effect in immature virions. These results show that incomplete maturation or binding of CD4 to mature HIV-1 virions induce a redistribution of HXB2 Env within the viral membrane and suggest a functional implication of the cluster conformation of Env during the pre-fusion reaction.

Seeking to know whether the intermolecular dynamics observed in HXB2 Env are strain-specific or instead it is a common behaviour shared across other HIV-1 strains, we performed similar photobleaching experiments with pseudoviruses bearing the R5-tropic, clinical isolate, JR-FL Env (Fig. 3) or the X4-tropic, laboratory-adapted, NL4-3 Env glycoprotein (Fig. 4) (tier 2 and tier 1A, respectively). In this case, pseudoviruses were produced labelling the HIV-1 Gag precursor with GFP but with unlabelled Env, to keep protein dynamics and their fusogenic activity under native conditions. HIV-1 mature or immature virions were incubated in presence or absence of soluble sCD4_D1-D4_ and the sample was later subjected to fixation and revealed by immunostaining (Fig. 3A). The anti-gp120 HIV-1 antibody b12 was used as primary antibody and an anti-human coupled to Alexa 633, was used as secondary antibody. It is worth noting that sample fixation was carried out before incubation with the b12 antibody to prevent any possible effect on Env distribution caused by the neutralizing antibody. Photobleaching was induced by continuous laser 633nm excitation of the Alexa 633 fluorophore on labelled virions. The resulting intensity traces and corresponding histograms were analyzed to obtain the average number of discrete photobleaching steps for each condition in both, HIV-1 JR-FL Env (Fig. 3B-C; Fig. S4) and HIV-1 NL4-3 Env viruses (Fig.4A-B; FigS5A). Interestingly, we could observe a similar tendency of the photobleaching dynamics in virions bearing JR-FL (Fig. 3C) or NL4-3 (Fig. 4B). In both cases, photobleaching events were significantly reduced, in a statistical manner, in immature viruses compared to mature virions, and the same effect was observed when exposing virions to soluble CD4. Of note, the number of photobleaching events in HIV-1 NL4-3 Env mature virions was higher (8.9±2.1) compared to JR-FL or HXB2 (3.8±1 and 5.1±1.1, respectively) pseudo-typed virions, which could reflect a difference in the relative number of Envs within the cluster, depending on the Env subtype. Photobleaching kinetics on JR-FL pseudo-typed virions turned out to be faster in mature virions compared to immature virions or in presence of its ligand, CD4 (Fig. 3D). This result was consistent with the photobleaching kinetics observed in mature vs immature HIV-1 NL4-3 Env virions, although the presence of sCD4 did not induce a significant delay in this case (Fig. 4C).

**Figure 3.**
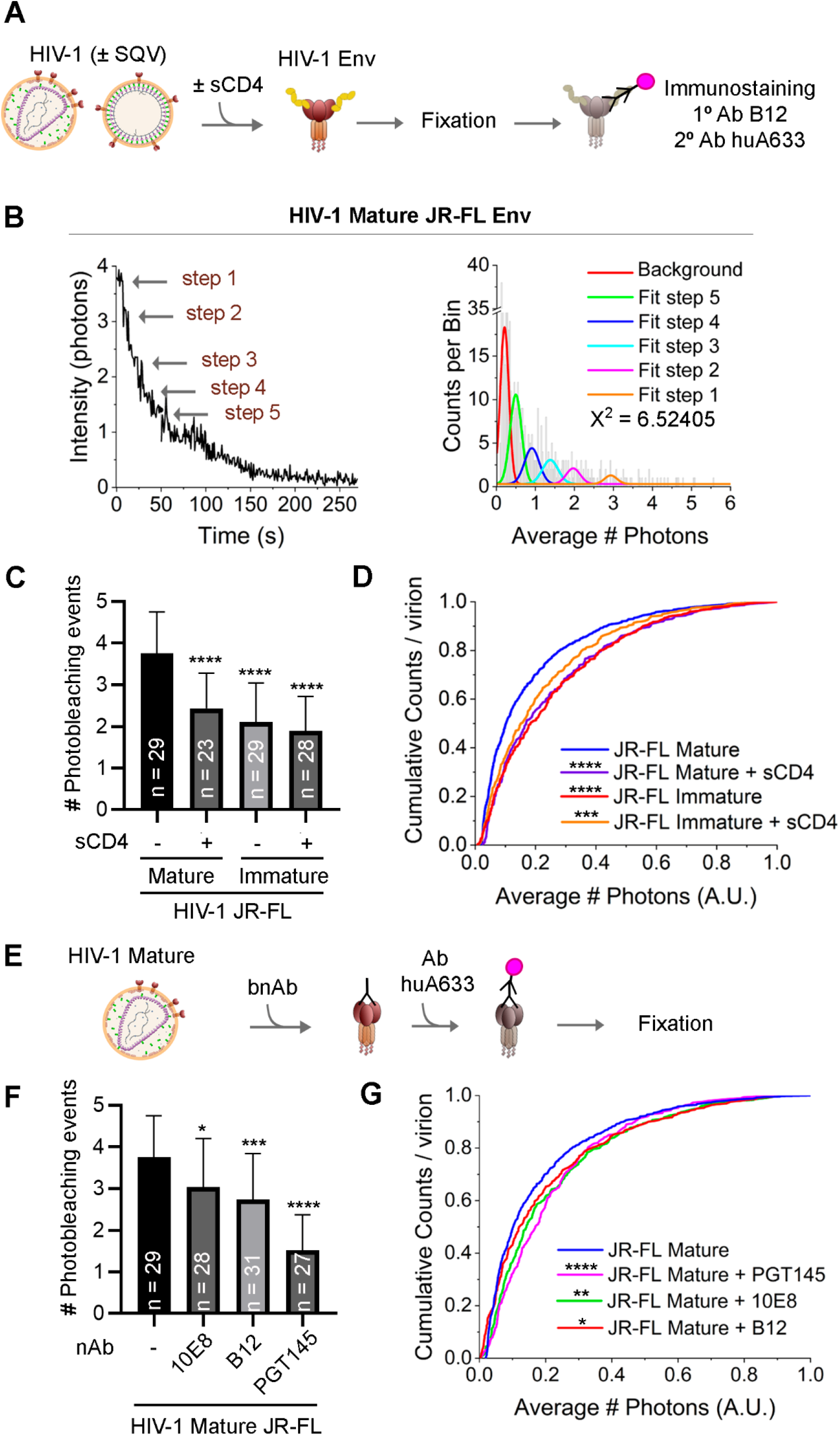
Intermolecular conformations of HIV-1 JR-FL Env are maturation-dependent and modulated by sCD4 and neutralizing antibodies. (A) Schematic representation of sample preparation for the single-molecule step photobleaching approach. Mature (-SQV) or immature (+SQV) HIV-1 particles were incubated in presence or absence of sCD4 and revealed by immunostaining with anti-gp120 b12 and anti-human A633 antibodies. Photobleaching of the Alexa 633 fluorophore labelling HXB2 V4-GFP_OPT_ was induced by continuous laser 633 nm excitation. (B) Representative intensity trace of HIV-1 mature JR-FL Env (left) and its corresponding population histogram (right) calculated from intensity traces. Data were fitted into a multi-Gaussian distribution model to estimate the number of photobleaching steps. Arrows point to single photobleaching steps detected. Χ^2^ values report the goodness of the fit. (C) Bar graph showing the mean and SD of the number of photobleaching events quantified from intensity traces. N of intensity traces is indicated on each bar per condition. Statistical significance was calculated using a one-way ANOVA and Sidak pos-hoc test comparing each condition to the mature HIV-1 JR-FL condition in absence of sCD4. **** < 0.0001; *** < 0.001; * < 0.05. (D) Cumulative distribution calculated from the mean intensity trace histograms from viral particles per condition. Statistical significance was calculated using a Kolmogorov-smirnov test comparing each condition to the mature HIV-1 JR-FL condition in absence of sCD4. **** < 0.0001; *** < 0.001; ** < 0.01; * < 0.05. (E) Schematic representation of sample preparation for the single-molecule step photobleaching approach. Mature HIV-1 particles were incubated in presence or absence of neutralizing antibodies 10E8, b12, PGT145 targeting Env and revealed by anti-human A633 antibodies. Photobleaching of the Alexa 633 fluorophore labelling HXB2 V4-GFP_OPT_ was induced by continuous laser 633 nm excitation. (F) Analysis as in (C). (G) Analysis as in (D).

**Figure 4.**
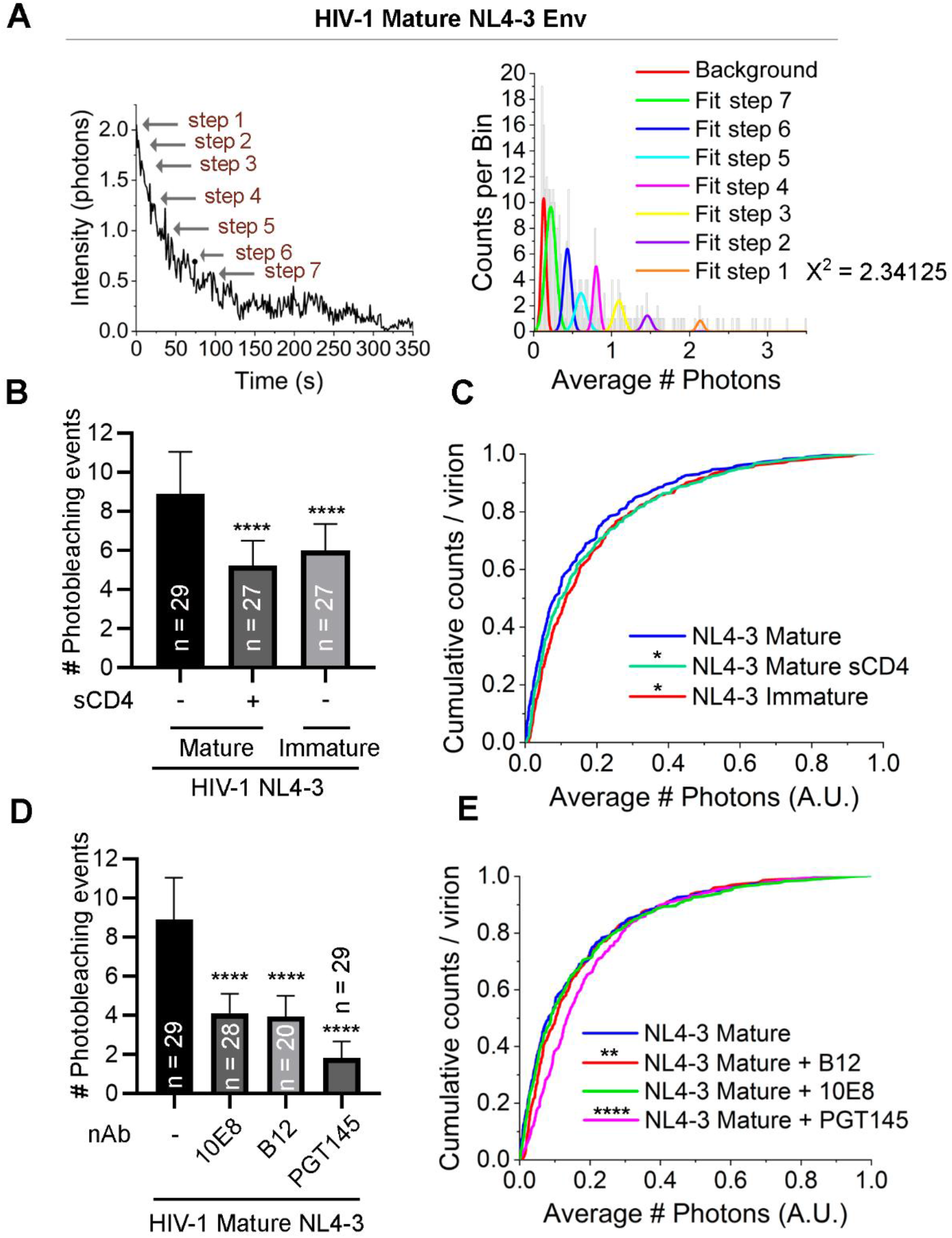
Intermolecular conformations of HIV-1 NL4-3 Env are maturation-dependent and modulated by sCD4 and neutralizing antibodies. (A) Representative intensity trace of HIV-1 mature NL4-3 Env (left) and its corresponding population histogram (right) calculated from intensity traces. Data were fitted into a multi-Gaussian distribution model to estimate the number of photobleaching steps. Χ^2^ values report the goodness of the fit. Arrows point to single photobleaching steps detected. (B) Bar graph showing the mean and SD of the number of photobleaching events quantified from intensity traces. N of intensity traces is indicated on each bar per condition. Statistical significance was calculated using a one-way ANOVA and Sidak pos-hoc test comparing each condition to the mature HIV-1 JR-FL condition in absence of sCD4. **** < 0.0001. (C) Cumulative distribution calculated from the mean intensity trace histograms. Statistical significance was calculated using a Kolmogorov-smirnov test comparing each condition to the mature HIV-1 NL4-3 condition in absence of sCD4. **** < 0.0001; ** < 0.01; * < 0.05. (D) Analysis as in (B) for HIV-1 NL4-3 Env mature virions incubated in absence or presence of neutralizing antibodies 10E8, b12 or PGT145. (E) Analysis as in (C) for HIV-1 NL4-3 Env mature virions incubated in absence or presence of neutralizing antibodies 10E8, b12 or PGT145.

With the aim to further characterize the functional aspect of the intermolecular conformations of Env, we investigated the effect of well-known HIV-1 broad neutralizing antibodies (bNAbs) on Env cluster formation. For this experiment, we selected three different antibodies recognizing separated regions of Env: PGT145, which recognizes the HIV-1 apex ^28^; B12, which binds to an overlapping region of gp120 with the site of CD4 attachment ^29,30^; and 10E8 which is directed against the membrane-proximal external region (MPER) ^31^. Moreover, these antibodies show high (PGT145 and 10E8) to moderate (b12) ability to neutralize different HIV-1 isolates ^32^. BNAbs were incubated with mature HIV-1 virions before sample fixation, and subsequently exposed to anti-human secondary antibodies coupled with Alexa 633 fluorophore (Fig. 3E). Strikingly, binding of bNAbs to HIV-1 JR-FL Env (Fig. 3F; Fig. S4B) or NL4-3 (Fig. 4D; Fig. S5B) induced a statistically significant reduction in the number of photobleaching events detected, with PGT145 showing the strongest effect, in both cases (1.5**±**0.8 for JR-FL and 1.8**±**0.8 for NL4-3). Incubation of mature HIV-1 JR-FL with bNAbs also caused a delay in photobleaching kinetics (Fig. 3G). A similar effect was observed when exposing HIV-1 NL4-3 Env virions to B12 and PGT145, but not to 10E8.

Overall, these experiments show that the cluster conformation of Env within the viral membrane depends on virion maturation. Furthermore, this conformation is impaired upon binding of Env to soluble CD4 and when mature virions are exposed to bNAbs targeting different epitopes of Env, suggesting a functional implication of intermolecular dynamics of Env during the pre-fusion reaction.

### Intermolecular Env clusters are destabilized during the prefusion reaction in live T cells

In order to investigate the intra- and intermolecular dynamics of HIV-1 Env in a physiological context, we studied the sequence of intra- and intermolecular transitions of HIV-1_Gag-GFP HXB2 V4-GFPopt_ virions labelled with donor (NbA488) and acceptor (NbA594) fluorophores when engaged to MT-4 T cells (Fig. 5A). We examined the time-resolved lifetimes and apparent FRET efficiencies that were simultaneously acquired at a time resolution of 3 s per FLIM image during 5 min (Fig. 5B). The three different E_app_ regimes previously described (low, E_app_ < 0.12; intermediate, 0.12 < E_app_ < 0.23 and high, E_app_ > 0.23) were taken as a reference to filter out each of the dwell times coming from individual E_app_ trajectories (Fig. 5B). The three dwell time distributions coming from at least 24 individual HIV-1 virions with a good signal to noise (between 100 and 1000 photons per pixel) were plotted as cumulative distribution functions (CDF) that, in turn, represent the average kinetics of each Env conformational states (Fig. 5C). The half lifetime of each one of the cumulative distributions provides quantitative information on the stability of each dynamic conformational state: a shorter CDF half lifetime implies a fast transition, and therefore, a very unstable Env conformational state, and a long CDF half lifetime translates instead in slow Env kinetics and stable Env conformational states. We first analyzed the CDF kinetics of mature HIV-1_Gag-GFP HXB2 V4-GFPopt_ particles *in vitro*. For the low E_app_ regime, corresponding to the open Env conformation, we observed a half lifetime of τ_(1/2)_ = 92 s. The intermediate E_app_ regime (0.12< E< 0.23) cumulative distribution kinetics, corresponding to the closed Env conformation gave a τ_(1/2)_ = 170 s. The long CDF lifetime of this particular closed Env conformation assumes a very stable and predominant state over the open conformation for unbound Env. In turn, the high FRET kinetic regime of Env cluster formation (E_app_ > 0.23) gave a τ_(1/2)_ = 22 s.

**Figure 5.**
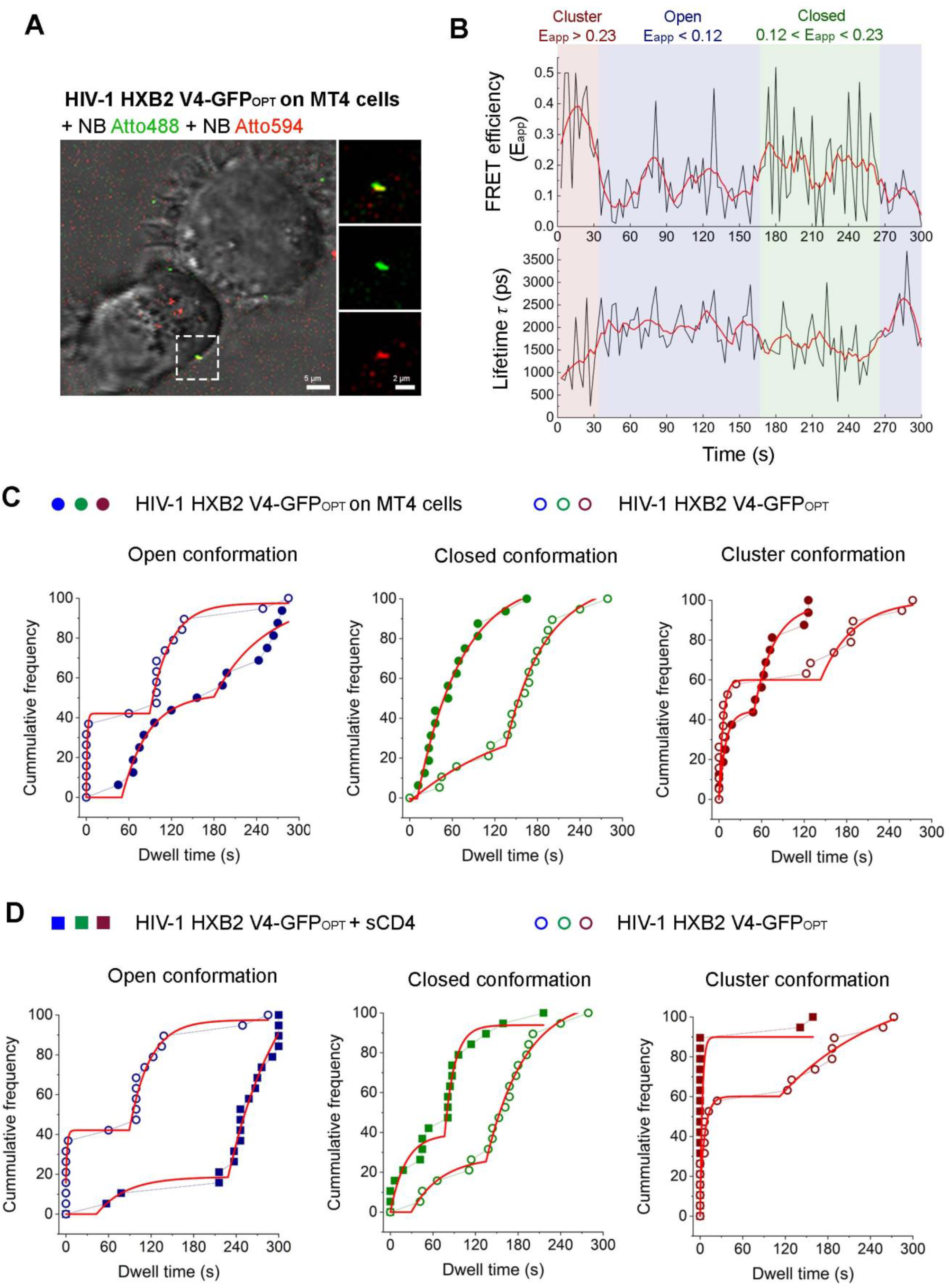
HIV-1 Env cluster is destabilized when engaged in the pre-fusion reaction in live T cells. (A) Micrograph showing mature HIV-1 HXB2 V4-GFP_OPT_ virions labelled with NbA488 (green) NbA594 (red) engaged in the pre-fusion reaction onto living MT4 T cells (phase contrast). Scale bar image on the left is 5 μm. Magnification of the region indicated by dashed lines is shown on the right. Viral particle showing colocalization between green and red channels is shown on the upper right panel. Middle and bottom right panels correspond to the same viral particle as observed in green and red channels, respectively. Scale bar magnification is 2 μm. (B) Graphs represent FRET efficiency (E_app_) (upper panel) and lifetime (in ps, bottom panel) traces over time. High FRET efficiency burst (E_app_ > 0.23), defining intermolecular interactions is depicted in red; intermediate FRET efficiency regime (0.12 < E_app_ < 0.23) assigned to closed Env conformations is depicted in green, and low FRET efficiency bursts (E_app_ < 0.12) reporting open Env conformations is depicted in blue. Note that hight FRET efficiency correlates with low lifetime values and *vice versa*. (C) Cumulative Distribution Functions (CDF) are plotted for E_app_ single traces obtained from at least (n = 20) HIV-1 HXB2 V4-GFP_OPT_ virions *in vitro* (open dots) and in presence of living T cells (solid dots). Each FRET regime determines the Env conformational state and kinetics. (D) Analysis as in (C) of HIV-1 HXB2 V4-GFP_OPT_ virions *in vitro* (open dots) and in presence of sCD4 (solid squares).

When the HIV-1_Gag-GFP HXB2 V4-GFPopt_ virions were engaged to living MT-4 T cells (n = 20), both the intramolecular and intermolecular Env landscape drastically changed compared to the previous condition with unliganded Env (Fig. 5C). A delayed and therefore more stable cumulative distribution kinetics was found for the populations corresponding to Env open conformation (τ_(1/2)_ = 152 s). The dynamic behaviour of the Env closed conformation in presence of T cells showed a half lifetime of τ_(1/2)_ = 58 s, which is faster and therefore less stable than in absence of T cells (Fig. 5C, middle chart); suggesting that Env might have already interacted with CD4 molecules exposed to the cell membrane of MT-4 T cells and hence, inducing an open conformation in the prefusion reaction ^5^. Finally, we observed that the Env cluster conformation when engaged to T cells was destabilized as compared to virions *in vitro* (τ_(1/2)_ = 12 s).

This generalized behaviour for the three Env conformational regimes defined above followed a similar tendency with the addition of sCD4_D1-D4_ (Fig. 5D). The open conformation was readily stabilized upon addition of the HIV-1 soluble receptor (τ_(1/2)_ = 253 s; Fig. 5D, left chart), as opposed to the closed conformation which was clearly destabilized (τ_(1/2)_ = 85 s; Fig. 5D, middle chart). Destabilization of the Env cluster upon addition of sCD4_D1-D4_ was more drastic compared to virions engaged to T cells (τ_(1/2)_ = 1 s, Fig. 5D, right chart), which might be the result of higher number of Env molecules binding to its receptor, due to the exposure of virions to saturating concentrations of sCD4_D1-D4._

We have thus shown that HIV-1 Env transitions towards an open conformation with longer CDF half lifetimes when engaged to T cells as compared to *in vitro* virions. This implies an overall increase in CDF half lifetime for Env open conformation of ~ 60 s (Δ_time_= τ_(1/2) (T cells)_ – τ_(1/2) (in vitro)_ = 152 s – 92 s = 60 s); concomitantly the CDF half lifetime for the Env closed conformation was shorter and more unstable with an overall decrease of ~ 1.86 min (Δ_time_= 58 s – 170 s = −112 s). Finally, Env cluster dissociation kinetics were also favored, giving rise to shorter CDF half lifetimes and more unstable Env intermolecular interactions with an overall decrease of 10 s (Δ_time_= 12 s – 22 s = −10 s). Overall, we have found that Env transitions between at open, closed and clustered conformational states. These Env dynamic states are shifted towards more stable Env open conformation, as opposed to the closed and cluster conformations in sCD4-bound virions or when primed to T cells, suggesting a potential dissociation of Env clusters into separate trimers upon engagement to CD4 on T cells.

### HIV-1 Env cluster disruption as a common mechanism for antibody neutralization

Next, we examined how the addition of inhibitory concentrations of different bNAbs affected the conformational dynamics of Env when engaged in the prefusion complex with living T cells (Fig. 6). HIV-1_Gag-GFP HXB2 V4-GFPopt_ viruses on the surface of MT4 T cells were exposed to inhibitory concentrations of PGT145, b12 and 10E8. These bNAbs do recognize different Env regions of vulnerability and show selective preferences towards specific Env conformations ^33^. PGT145 has been previously reported to recognize the quaternary structure of the Env apex trimer, thus, stabilizing a closed conformation of HIV-1 Env ^28^. B12, as opposed to CD4, is unable to bind the closed conformation of Env, although upon binding, b12 prevents reversion back to the closed state ^29^, thus stabilizing an intermediate/open conformation ^30^. Finally, 10E8 targets a quaternary epitope including lipid and MPER contacts ^31^ stabilizing an open conformation ^25^.

**Figure 6.**
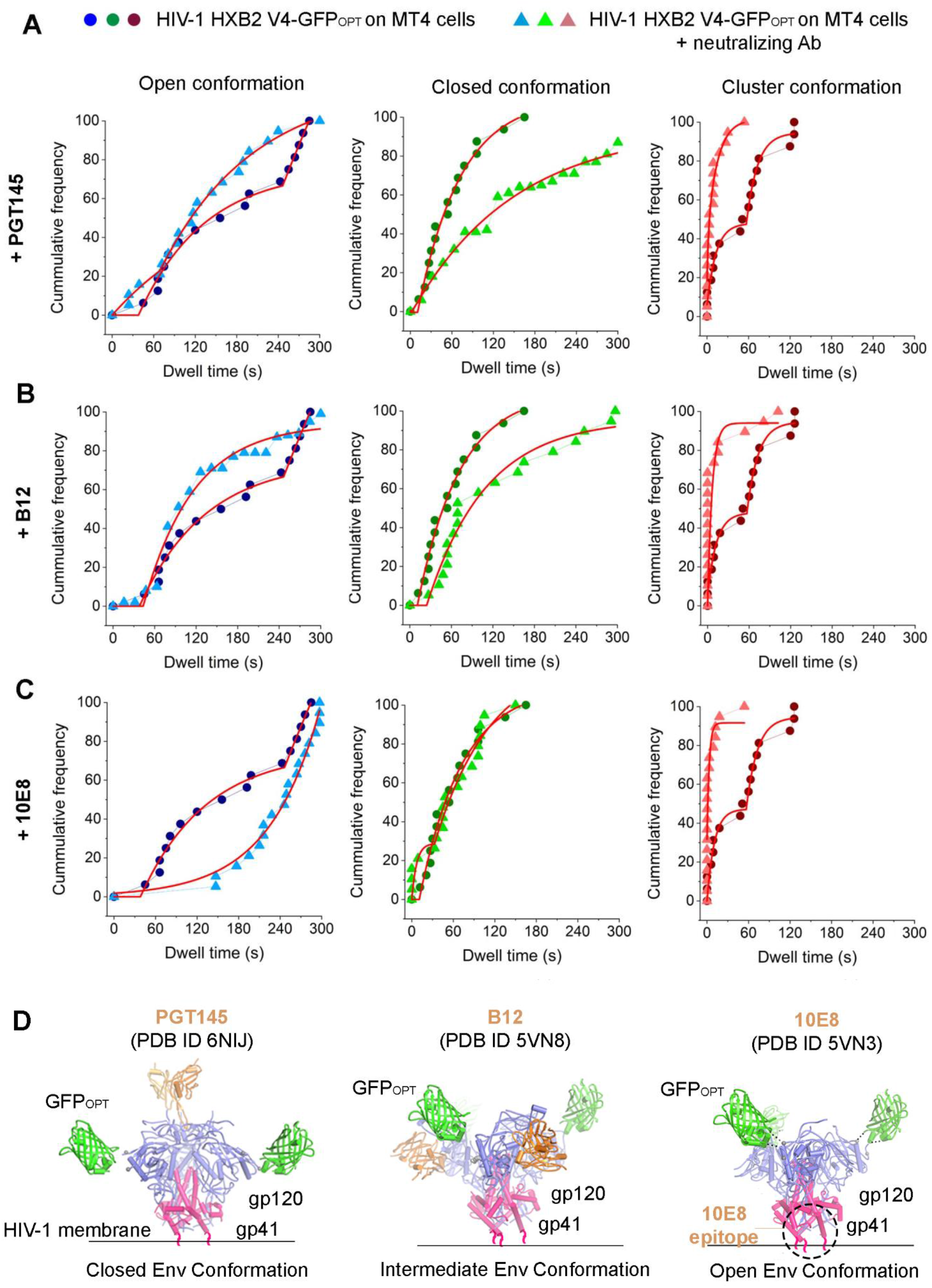
HIV-1 Env cluster is destabilized when exposing virions to neutralizing antibodies in live T cells. Cumulative Distribution Functions (CDF) are plotted for FRET efficiency traces obtained from at least (n = 20) HIV-1 HXB2 V4-GFP_OPT_ virions in presence of living T cells without (solid dots) or with (solid triangles) neutralizing antibodies PGT145 (A), b12 (B) and 10E8 (C). Each FRET regime (high, E_app_ > 0.23 in red; intermediate, 0.12 < E_app_ < 0.23, in green; low, E_app_ < 0.12, in blue) determines the Env conformational state and kinetics. (D) The HIV-1 Env structures representing closed Env associated with PGT145 (PBD ID 6NIJ, left), b12 (PBD ID 5VN8, middle) or 10E8 (PBD ID 5VN3, right) were modified to illustrate the labeling of the V4 loop with GFP_OPT_.

Our smFRET data indicate that PGT145 exhibited a tendency to destabilize the open Env conformation triggered by Env binding to the CD4 T cell receptor (from τ_(1/2)_= 152 s to τ_(1/2)_ = 118 s) and, consequently, reversing the CDF half lifetime towards a closed Env conformation (from τ_(1/2)_ = 58 s to τ_(1/2)_ = 120 s) (Fig. 6A, left and middle chart). We also observed a destabilization of the Env open conformation when virions were incubated with T cells in presence of the b12 neutralizing antibody (from τ_(1/2)_= 152 s to τ_(1/2)_ = 1 s). Concomitantly, b12 induced a stabilization of the closed Env conformation (from τ_(1/2)_ = 58 s to τ_(1/2)_ = 92 s), although less efficiently than the apex-directed PGT145 antibody (Fig. 6B, left and middle chart). In turn, the anti-MPER antibody, 10E8, induced a strong stabilization of the open Env conformation (from τ_(1/2)_ = 152 s to τ_(1/2)_ = 250 s) and consistently, the closed conformation kinetics were very similar in absence or in presence of the antibody (from τ_(1/2)_ = 58 s, to τ_(1/2)_ = 58.2 s) (Fig. 6C, left and middle chart). Interestingly, the three antibodies tested induced a drastic destabilization of the Env cluster (from τ_(1/2)_= 12 s to τ_(1/2)_ = 3 s, for PGT145; to τ_(1/2)_= 1 s, b12; to τ_(1/2)_ = 0.8 s, 10E8) (Fig. 6A-C, right chart).

In light of these results, a structural model summarizing the intramolecular conformations of HXB2 V4-GFP_OPT_ Env observed for each neutralizing antibody is shown in Fig. 6D. In our system, a closed Env conformation was stabilized when incubating HIV-1 virions engaged to T cells in presence PGT145, whereas the b12 antibody favours an intermediate intramolecular conformation of Env and, instead, a stable open conformation was observed upon 10E8 addition. Furthermore, these results show that bNAbs, which are known to stabilize intramolecular conformations of Env, strongly impair intermolecular dynamics of Env by destabilizing and dissociating the Env cluster during the pre-fusion reaction of HIV-1 virions on T cells. Therefore, these results suggest a common mechanism of Env cluster disruption by bNAbs even though each one of them binds to different Env regions. Moreover, these data also point to Env cluster dissociation as an effective and potentially common strategy to inhibit HIV-1 fusion with T cells.

## Discussion

We have found a previously underappreciated mechanism involving Env intermolecular dynamics that might be crucial during the prefusion reaction. We have quantitatively shown how Env intermolecular interactions are reduced when primed to live MT-4 T cells. Furthermore, we have shown that three different families of bNAbs, targeting different Env epitopes (PGT145 targets the apex, b12 the CD4-binding region and 10E8 the membrane proximal external region (MPER) of Env) ^33^, disrupt Env clusters both *in vitro* (Fig. 3–4) and when engaged to live T cells (Fig. 6). Moreover, we have observed that cluster formation and dissociation by CD4- or bNAb-binding is a common mechanism shared across laboratory-adapted HXB2 and NL4-3 strains, and the clinical isolate JR-FL strain. In addition, our FRET-FLIM imaging system allowed us to reconcile dynamics of intra- and intermolecular interactions of Env, which dynamically transits between open and closed conformations and cluster association and dissociation in mature, unligated HIV-1 virions. These data were further validated with built in controls within the same experiments which gave us concomitant closed Env conformation stabilization upon Env binding to PGT145, an intermediate Env conformation in case of b12 and open Env conformation stabilization when bound to 10E8 or CD4 (Fig. 7).

**Figure 7.**
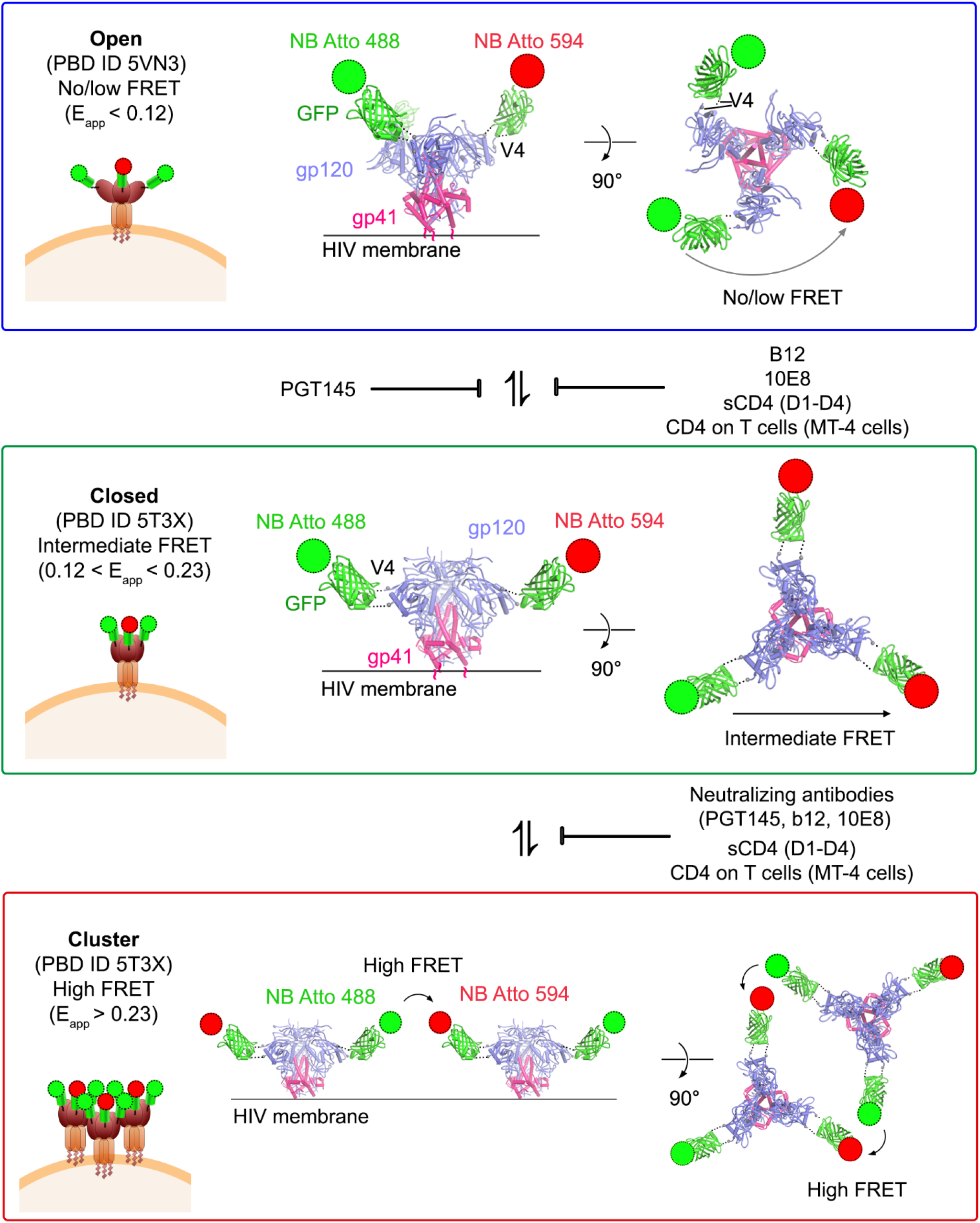
HIV-1 Env transitions between intra- and intermolecular conformations are disrupted by CD4-binding and bNAbs. Schematic representation of the transitions between different intra-(open, showing low/no FRET, upper panel; closed, showing moderate FRET, middle panel) and intermolecular (cluster, showing high FRET bottom panel) env conformations observed by FRET-FLIM. Receptor binding (sCD4_D1-D2_ or CD4 on T cells) and binding of b12 and 10E8 bNAbs stabilize and intermediate or open conformation, whereas PGT145 favours a closed conformation. Engagement of Env with CD4 or with any of bNAbs tested increases the distance between Env molecules, inducing cluster dissociation.

### Detection of intra- and intermolecular HIV-1 Env dynamics

The experimental design in this study has been crucial to evaluate intra- and intermolecular interactions of Env in native virions *in vitro* and engaged to T cells. Key parameters include the time-scale of acquisitions, the labelling strategy and resolution of dynamic interactions.

Previous results on Env intramolecular dynamics were recovered employing single molecule FRET combined with Total Internal Reflection Microscopy (TIRF) ^8,9,21^. This technology combined with dually labeled Env molecules using short peptides introduced into different gp120 loops provided an outstanding platform to evaluate V1 loop conformational changes ^9^. The time-acquisition for their single molecule FRET experiments was restricted to ~10 s (with 25 frames per second) prior to bleaching of the fluorophores. Although with a great time-resolution, this technique would fail to detect long-time lapse Env dynamics (in the range of seconds and minutes). Importantly, the high power laser needed to collect enough photons for the analysis would induce virus phototoxicity ^34^, which in turn might affect Env conformational dynamics. Of note, our two photon FRET-FLIM data were acquired for 5 min with a FLIM time resolution of 3 s, with minimal photobleaching. Moreover, according to our own data presented here, and to a number of biophysical studies ^3,12^ a longer time-scale might be crucial to detect Env intermolecular conformational dynamics given the slow Env diffusion coefficient in mature HIV-1 particles (D = 0.002 μm2/sec in mature particles ^3^. In this sense, the excitation source utilized in this work has been key to visualize HIV-1 Env dynamics engaged to living T cells. Two photon excitation provides high three-dimensional contrast and resolution without the need for optical filters (i.e. pinhole or notch filters) in the detection path. This, combined with digital photon counting (HyD) descanned detectors situated very close to a high numerical aperture objective, gave a high signal-to-noise ratio. Since two photon excitation is naturally confocal ^35^, only the viruses engaged in the MT-4 T cells were imaged and all emission photons gave a valuable signal, with reduced phototoxicity, allowing longer acquisitions times whilst conserving resolution and high contrast. Lastly, two photon excitation also provided a localized excitation were all emission photons constitute a useful signal that contributed to our rapid FLIM acquisitions.

The labeling strategy when performing FRET experiments is also crucial. A labeling approach strictly circumscribed to Env could be a restraint. In this scenario, one could not guarantee that all particles analyzed could be bona fide HIV-1 virions with their corresponding capsids. Here, we tagged the HIV-1 capsid (Gag-GFP) and the gp120 V4 domain of Env with a super-folding GFP (GFP_OPT_) ^14^. Even if all emitted photons from these labels are green, lifetime imaging permitted discrimination of mature versus immature viruses. On top of that, we employed labelled nanobodies ^5^ that specifically bind to GFP (in our system, only GFP_OPT_ is accessible to nanobodies) that could be accurately detected given the difference in lifetime signatures between NbA488 and GFP (Fig. 1–4). At least 5 amino acid residues linking GFP_OPT_ with the V4 and V1 loops of gp120 allow free, or random rotation of the fluorophores, thus, minimizing problems related to the relative dipole-dipole orientation (and here, one could approximate K^2^ = 2/3). Under these circumstances, FRET interpretation is restricted to protein folding and protein-protein interactions. (Fig. 7 and Fig. S5).

Choosing the label location within the Env glycoprotein is not trivial. To resolve both, intra- and intermolecular interactions, the V4 loop of the gp120 was selected, as the side location of this residue facilitates FRET to occur between different Env trimers. Also, in our system, all Env proteins within HIV-1 virions were labelled with GFP_OPT,_ increasing the presence of multiple acceptors per donor molecule in conditions of Env cluster formation and thus, facilitating an increased energy transfer and an additional shortening of the donor lifetime ^36^.

In previous reports, Env cluster characterization has been performed applying techniques such as STED ^3,12,37^, 3D STORM ^38^ or Cryo-ET ^39,40^. While these techniques offer incredibly high spatial resolution, they are unable to resolve Env conformational dynamics. Here, we employed a FRET-FLIM approach as a molecular ruler for the study of different Env conformation transitions on the nanoscale (∼2–10 nm). This technique allowed us to detect subtle modulations of Env trimer interactions in real time when binding to neutralizing antibodies or upon CD4-receptor binding of HIV-1 virions onto T cells, that were previously unappreciated ^12,37^.

Single Step Photobleaching, has previously been applied to HIV-1 Env *in vitro* ^41^. This approach was able to produce quantitative results on the number of soluble CD4 molecules per Env SOSIP.664 purified and immobilized in a glass coverslip. Here, we have taken this approach one step further by employing single step photobleaching on native HIV-1 virions. Even if Single Step Photobleaching is unable to produce time-resolved data and information on the Env intramolecular dynamics; we could resolve Env clusters in mature particles for the three Env tested, reinforcing the idea of the functional relevance of Env clusters as a pre-requisite to start the fusion reaction.

### Env conformation dynamics are modulated during HIV-1 prefusion reaction and are disrupted by bNAbs

In previous studies, HIV-1 Env has been described to exist in at least three intramolecular conformational states by using smFRET approaches ^7–9,21^. Unligated Env dynamically transits between different intramolecular conformations, although with a preference to pre-triggered, closed conformation (state 1). Binding to CD4 would first induce an asymmetric opening of Env (state 2) that ultimately leads to binding to its coreceptor CXCR4 or CCR5 (state 3) for subsequent fusion with the host membrane. Ensemble analysis of E_app_ in our system, resolved two different intramolecular conformations. We designed them as open and closed conformations. Open conformation showed the lowest FRET efficiency and was stabilized upon binding to sCD4_D1-D4_ in mature and immature virions. The concentration used in this assay (10 μg/mL) has been associated to three-CD4-bound conformation ^41^, hence we might relate this conformation with previously described as state 3. Closed conformation was characterized in our approach by moderate FRET efficiency regimes and was predominant in unligated mature virions. We further confirm this hypothesis when analyzing E_app_ time-resolved tracks from mature HIV-1 particles primed to T cells in presence of the PGT145 bNAb, which is known to have a preference for the closed Env conformation ^28^ and induced stabilization of the moderate FRET efficiency population in our assays. Therefore, the closed env conformation in our system could relate to previously assigned as state 1 conformation. Intermediate opening of Env was not clearly resolved by our ensemble E_app_ analysis. However, we observed subtle differences in the degree of stabilization of the open conformation when comparing E_app_ kinetics in mature HIV-1 in absence or presence of T cells or in absence or presence of the bNAbs b12 or 10E8, suggesting the existence of intermediate open states with different degrees of stability as previously observed ^8,25^.

When comparing multiparameter plots of E_app_ and lifetime of mature vs immature virions, we observed that high FRET efficiency regimes corresponded to intermolecular interactions of Env, as they were only observed in mature virions labelled at the V4 loop of gp120. When taking a closer look at E_app_ time-resolved tracks from mature HIV-1 particles *in vitro*, and also primed to T cells, the transition towards Env clusters always occurred via the Env closed conformation, and never from an Env open conformation (Fig. 7). We also observed dissociation of intermolecular Env interactions when HIV-1 virions were primed with T cells or in presence of sCD4_D1-D4_. These observations suggest that Env cluster formation requires Env to adopt a closed conformation, and its dissociation is triggered by receptor binding.

Using a single-photobleaching approach, we were able to see statistically relevant differences in both the number of steps and the photobleaching kinetics for three HIV-1 virions pseudotyped with NL4-3, HXB2 and JR-FL. We found that at least three Envs are incorporated in clusters for HIV-1 mature viruses decorated with NL4.3 Env (~9 photobleaching steps) and at least two in HXB2 and JR-FL (~5 photobleaching steps in each). Provided that most likely not all Env trimers were fully labeled, this estimation would count for the minimal amount of Env particles present in each cluster. This approach clearly shows two things, first, mature particles irrespective of Env (tier 1A, 1B and 2) tend to form clusters and second, receptor binding in all cases disrupts these clusters. In all cases, the addition of ligands (either sCD4_D1-D4_ or bNAbs) disrupted the Env cluster; only seen in mature HIV-1 virions.

Overall, our data clearly shows that Env cluster association and dissociation is a key factor in mature HIV-1 particles and play a fundamental role during immune evasion. Moreover, the mechanism of masking different Env regions by neutralizing antibodies might also be related to the intermolecular dynamics of Env and immune evasion. Taken together, this work suggests that destabilizing Env clusters could represent a common strategy to arrest and inhibit viral fusion machines.

## Acknowledgements

We thank Zene Matsuda for the kind gift of HIV-1 Env labeled plasmids. We thank Leica Microsystems for technological support. We thank the Padilla-Parra lab for valuable discussions and criticism of the paper. This work has been supported by the European Research Council (ERC-2019-CoG-863869 FUSION to S.P-P.) and the Wellcome Trust Core Award (203141).

## Author Contributions

Conceptualization, S.P.-P.; methodology, I.C.-A., T.M. and S.P.-P.; validation, I.C.-A. and S.P.-P.; formal analysis, I.C.-A. T.M and S.P-P; investigation, I.C.-A, T.M. and S.P.-P.; resources, S.P.-P.; data curation, I.C.-A. and S.P.-P.; writing’original draft preparation, S.P.-P.; writing’review and editing, I.C.-A., T.M. and S.P.-P.; visualization, I.C.-A. T.M and S.P-P; supervision, S.P.-P.; project administration, S.P.-P.; funding acquisition, S.P.-P. All authors have read and agreed to the published version of the manuscript;

## Material and Methods

### Cell Culture

Lenti-X™ 293T cells (Takara Bio, Clontech, Saint Germain en Laye, France) were grown using complete Dulbecco’s Modified Eagle Medium F-12 (DMEM F-12) (Thermo Fisher Waltham, MA, USA), supplemented with 10% fetal bovine serum (FBS), 1% penicillin-streptomycin (PS), and 1% L-glutamine. MT4 T (provided by Alex Compton, NCI Center for Cancer Research, Frederick, MD, USA) were cultured in RPMI 1640 medium containing 10% FBS, 1% PS and 1% L-glutamine. Cells were maintained in a 37 °C incubator supplied with 5% CO_2_. For experiments, MT4 T cells were cultured in PBS 1x buffer containing 2% FBS and 15 mM HEPES.

### Reagents and antibodies

Nanoboosters (Chromotek, Germany) targeting the GFP_OPT_ of labeled Env GFP_Booster_Atto488 and/or RFP_Booster_Atto594 (ChromoTek GmbH, Planegg, Germany) were used in 1:200 final concentration. Human soluble CD4 recombinant protein (sCD4_D1-D4_; Cat.No:4615, NIH AIDS reagent program) and broad neutralizing antibodies: anti-HIV-1 gp120 monoclonal, PGT145 (Cat. No: 12703, NIH AIDS reagent program); anti-HIV-1 gp41 monoclonal, 10E8 (Cat. No: 12294, NIH AIDS reagent program) and anti-HIV-1 gp120 monoclonal, b12 (Cat. No: AB011, Polymun Scientific, Klosterneuburg, Austria), were used in FRET-FLIM and SMPB experiments.

### Plasmid constructs

The pR8ΔEnv plasmid (encoding HIV-1 genome harbouring a deletion within Env), pcRev, NL4-3 Gag-iGFPΔEnv were kindly provided by Greg Melikyan (Emory University, Atlanta, GA, USA). The plasmids encoding the HXB2 gp120 V4 and V1 labeled with GFP_OPT_ were a kind gift from Zene Matsuda (Institute of Biophysics, Chinese Academy of Sciences, China). Plasmid encoding the JR-FL Env was a kind gift from James Binley (Torrey Pines Institute for Molecular Studies) and NL4-3 Env coding plasmid was provided by Dr Alex Compton.

### Virus production

Gag-GFP-containing, HXB2 V4-GFP_OPT_ and HXB2 V1-GFP_OPT_ pseudotyped viral particles were produced via transfection of 60-70% confluent Lenti-X™ 293T cells seeded in T175 flasks. DNA plasmids were transfected into Lenti-X™ 293T cells using GeneJuice^®^ (Novagen, Waltford, UK) according to manufacturer’s protocol. Specifically, cells were transfected with 2 μg pR8ΔEnv, 1 μg pcRev, 3 μg of NL4-3 Gag-iGFPΔEnv and 3μg of the appropriate viral envelope. All transfection mixtures were then added to cells supplemented with complete DMEM F12, upon which time they were incubated in a 37 °C, 5% CO_2_ incubator. 12 hours post-transfection, the medium was replaced with fresh, phenol-red free, complete DMEM F12 after washing with PBS. In the case of immature HIV-1 pseudovirus production, complete DMEM F-12 was supplemented with 300 nM of the HIV-1 protease inhibitor Saquinavir mesylate (Sigma-Aldrich, St. Louis, MO, USA). 72 h post-transfection, the supernatant containing virus particles was harvested and filtered with a 0.45 μm syringe filter (Sartorius Stedim Biotech). Filtered viral supernatants were concentrated 100 times using Lenti-X™ Concentrator (Takara Bio, Clontech, Saint Germain en Laye, France) and resuspended in phenol red-free medium, FluoroBrite DMEM (Thermo Fisher, Waltham, MA, USA), aliquoted and stored at −80 °C.

### Sample preparation

HIV-1 viruses pseudotyped with labelled HXB2 Env and harbouring Gag-GFP were diluted in PBS 1x, 2% FBS buffer and plated onto a micro-slide (Cat.No: 81826, Ibidi, Gräfelfing, Germany) by centrifugation at 2100 g, 4°C during 20 min. Unbound viruses were removed and media replaced by diluted nanoboosters (NbA488 and/or Nb594) in presence or absence of 10 μg/mL sCD4_D1-D4_ or 2 μM concentration of bNAbs in PBS 1x, 2% FBS at 20 μl final volume. The sample was then incubated for 1h at room temperature (RT) before image acquisition. In case of sample preparation for SMPB experiments, labelling of HXB2 V4-GFP was done after incubation for 1h at RT in absence or presence of sCD4_D1-D4_ followed by sample fixation using 4% PFA in PBS 1x for 15 min at RT. Virions were then incubated with 1:200 diluted NbA594 for 1h at RT. Labelling of HIV-1 viruses pseudotyped with JR-FL or NL4-3, and harbouring Gag-GFP was done after incubation for 1h at RT in absence or presence of sCD4_D1-D4_ or bNAbs at the concentrations indicated above. The sample was then fixed using 4% PFA in PBS 1x for 15 min at RT. In case of virions incubated ± sCD4_D1-D4_, the fixed sample was incubated with 1:500 dilution of b12 antibody for 1h at RT. Goat anti-human coupled to Alexa 633 fluorophore (Invitrogen) was used as secondary antibody diluted 1:500 and incubated for 30 min at RT. In case of virions incubated with bNAbs, the fixed sample was incubated with 1:500 dilution of goat anti-human coupled to Alexa 633 fluorophore (Invitrogen) and incubated for 30 min at RT. Antibodies for immunostaining were diluted in PBS 1x in presence of 2% FBS to prevent unspecific binding and incubations were done in the dark to preserve the fluorophore staining.

### Single Virus Tracking

MT-4 T cells were added onto surface-bound viruses in a final volume of 20 μl. Cells were spun for 10 min at 600 g in a refrigerated centrifuge so that the HIV-1 particles could engage with the cells, without initiating the prefusion reaction. The observation micro-slides were then put under the microscope and the cold medium immediately replaced by medium at RT right at the moment when we started the imaging acquisition procedure. We employed the two photon SP8 X SMD DIVE FALCON confocal microscope (also described below) equipped with a dark incubator chamber for the frame.

Virus tracking was performed with both ImageJ plugin Spot tracker and the 64-bit software module from Imaris (BitPlane, Zurich, Switzerland), using an auto-regressive algorithm. Tracking provided quantitative information regarding the mean fluorescence intensities of the HIV-1 HXB2-GFP_OPT_ with NbA488 (donor) collected in the non-descanned HyD1, the sensitized emission of the donor in the presence and absence of the acceptor HIV-1 HXB2-GFP_OPT_ NbA488 (donor) and Nb594 (acceptor) was recovered in the second non-descanned HyD2. Tracking of individual particles both in vitro and when engaged in MT-4 T cells also provided FLIM values for the donor, particle’s instantaneous velocity, trajectory and the mean square displacements (MSD).

### Fluorescence lifetime imaging microscopy (FLIM)

In vitro HIV-1 labeled virions (mature and immature HIV-1 HXB2-GFP_OPT_ NbA488 (donor)-Nb594 (acceptor)) and live MT-4 T cells exposed to HIV-1 particles were imaged using a DIVE SP8–X-SMD FALCON Leica microscope, Leica Microsystems (Manheim, Germany). Both, HIV-1 virions and MT-4 T cells of interest were selected under a 100x/1.4 oil immersion objective corrected for infra-red light (IR). HIV-1 labeled virions were excited using a two-photon femtosecond pulsed laser tuned at 950 nm and 80 Mhz. The FALCON module was coupled with single photon counting electronics for rapid FLIM (Leica Microsystems) with virtual gating set at 97 ps. Green emission photons were subsequently detected by three hybrid non-descanned external detectors in photon counting mode with emission filters set at 500-550 nm, 600-650 nm and a long pass starting at 700 nm for the third HyD detector. Stacks of 100 images of time-resolved data acquired at 1-3 sec each for 5 minutes were acquired for all experiments. HIV-1 HXB2-GFP_OPT_ Nb594 (acceptor) particles were tested to avoid the possibility of cross-excitation of the acceptor (Nb594) with the two-photon laser tuned at 950 nm. No photons were detected in the acceptor channel with the power set at 10% of the laser power (Spectra Physics, UK). Leica software (LAS X) was employed to produce the phasor plots (Leica Microsystems, Mannheim, Germany). ImageJ (https://imagej.nih.gov/ij/) and Originlab (Northhampton, USA) were employed to produce the multiparameter two dimensional graphs and probability kernel maps comparing the apparent FRET efficiency (calculated as described below) with average lifetime data (given in picoseconds per pixel) and recovered with LAS X and previously treated with ImageJ to remove the noise.

Both, photon counting images for the donor (HyD1) and the sensitized FRET emission and FLIM micrographs simultaneously acquired by the two photon SP8 DIVE FALCON system using the same microscopy settings were background subtracted, to get rid of the scatter photons and white noise recovered by each HyD channel. After this, each single virus was profiled utilizing a mask that only contained the signal coming from each individual particle (in vitro or engaged in non-labeled T cells) and non-attributed-numbers for the background. Both, the time-resolved intensity and average lifetime values for each channel were obtained together with the average intensity values (in photons and not grey values). The average number of photons and average lifetime per channel for each profiled virus was obtained (n > 100 particles for in vitro and n > 65 for live cell imaging) and plotted as a multiparameter plot and phasor plot for each condition. Individual traces for each condition were also recovered following this procedure.

### FRET and FLIM image analysis

FRET, is a nonradioactive, dipole-dipole coupling process where the energy is transferred from the excited donor fluorophore to the acceptor fluorophore when the distance and orientation of both dipoles are the right ones (typically within 10 nm and a random orientation). The excitation of the donor fluorophore induces a sensitized emission from the acceptor concomitantly quenching the fluorescence of the donor. This process, in the absence of acceptor would not occur. The HXB2 Env was fused to genetically encoded GFP_OPT_ that in turn was labeled by nanoboosters (NbA488, playing the role of the donor and Nb594, playing the role of the acceptor), and the molecular dynamics of Env in question was then inferred by FRET between the fluorophores. The efficiency with which Förster-type energy transfer occurs in given by the next equations:

The FRET efficiency (E) can be calculated as the proportion of photons absorbed in the donor versus the excitation transferred to the acceptor:

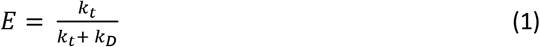

Where k_D_ is the sum of all relaxation pathways and k_T_ the transfer rate.

Experimentally we calculated E_app_ pixel by pixel utilizing the next equation

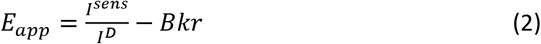

The signal of the laser pulse and delayed photon arrivals were rapidly digitalized at high speed with a temporal resolution per channel of 97 ps, allowing very rapid FLIM acquisitions (1-3 sec per FLIM image). Pixel by pixel images with their corresponding background subtracted average lifetime images where directly provided by the Leica software LAS X with the FALCON module. Single Photons coming from the donor/s (HIV-1 HXB2 V4 or V1-GFP_OPT_ labelled with NbA488 (donor) in the presence and absence of NbA594 (acceptor)) were detected in the non-descanned HyD detector. FLIM analysis was performed applying the non-fitting phasor plot approach fully integrated in the LAS X software.

### Intramolecular and intermolecular dynamics model

The average time-resolved lifetimes and apparent FRET efficiencies that were simultaneously acquired at a time resolution of 3 sec per FLIM image during 5 minutes, coming from individual HIV-1 viruses were filtered using the next criteria (for each condition). The three different E_app_ regimes previously described (high, E_app_ > 0.23, intermediate 0.12 < E_app_ < 0.23 and low E_app_ < 0.12) were taken as a reference to select each of the dwell times coming from individual E_app_ trajectories. The three dwell time distributions coming from at least 20 individual HIV-1 viruses per condition; were plotted as cumulative distribution functions (CDF). Only HIV-1 particles with a good signal to noise (between 100 and 1000 photons per pixel) were selected for all conditions. The t(1/2) was recovered for each CDF curves.

### Single Step Photobleaching acquisition and analysis

In vitro HIV-1 virions (mature and immature HIV-1 HXB2 V4-GFP_OPT_, HIV-1 NL4.3 Env and HIV-1 JR-FL Env). were selected under a 100x/1.4 oil immersion objective corrected for infra-red light (IR). HIV-1 double labeled virions (containing Gag-GFP and labeled Env) were excited using either and HeNe laser tuned at 591 nm or an Argon laser tuned at 644 nm at high power. The SP8 system (described above) was coupled with single photon counting HyD detectors (Leica Microsystems). Red emission photons were subsequently detected by these hybrid detectors in photon counting mode with emission filters either set at 600-650 nm or 660 750 nm for Atto 594 and 644 respectively. ImageJ (https://imagej.nih.gov/ij/) and Originlab (Northhampton, USA) were employed to produce the single step photobleaching graphs and histograms All data was previously treated with ImageJ to remove the noise.

### Structural Models and Analysis

A model for intra and intermolecular dynamics of the HXB2 Env labeled with GFP_OPT_ and nanoboosters was built using the following structures: ligand-free HIV-1 Env mimic (BG505 SOSIP.664) (PDB: 4ZMJ), HIV-1 Env mimic (B41 SOSIP.664) in complex with the ectodomain of CD4 (PDB: 5VN3), HIV-1 Env mimic (B41 SOSIP.664). Models were generated in Chimera, Coot, and Pymol (https://pymol.org).

### Statistics

For FRET-FLIM analyses, calculation of multiparameter 2D kernel density plots and t(1/2) of CDF from E_app_ traces was performed using Originlab software (Northhampton, USA). Statistical analyses comparing the mean of the number of photobleaching events between conditions was done applying a one-way ANOVA and Sidak’s multiple comparision test (significance p < 0.05) calculated in GraphPad Prism 8 software (California, USA). Statistical comparison between photobleaching kinetics was performed applying a Kolmogorov-Smirnov test (significance p < 0.05). Resulting histograms from photobleaching intensity traces were fitted to a multi-Gaussian model and the goodness of the fit was judged by X^2^ values closest to 1. These analyses were performed using Originlab software.

---

**Figure S1.**
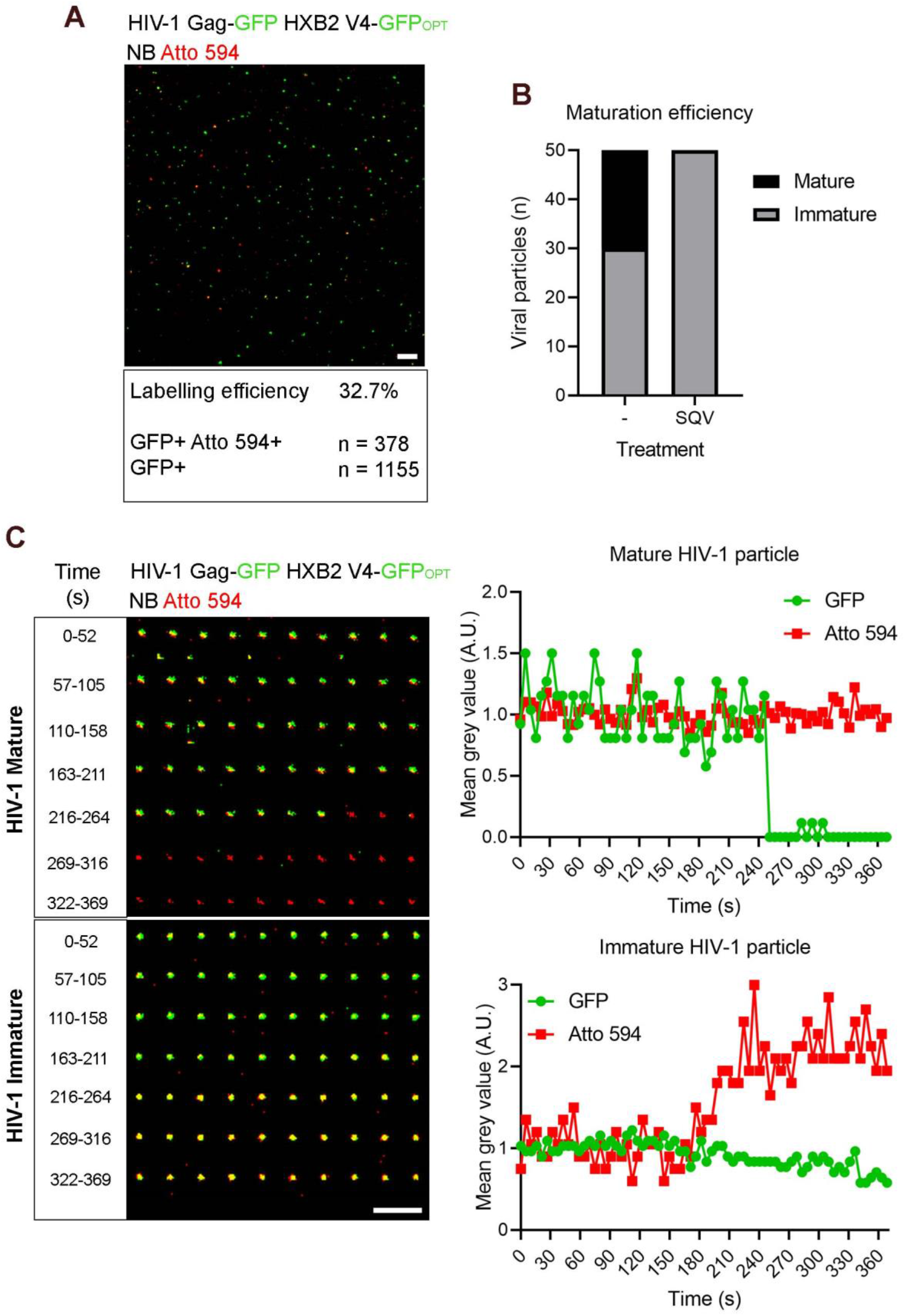
Efficiency of labelling and maturation of HXB2 V4-GFP_OPT_ virions. (A) Micrograph showing the labelling efficiency of HIV-1 pseudoviruses expressing Gag-GFP and HXB2 V4-GFP_OPT_ (green) labelled with Atto 594 (red). Colocalisation (GFP+ Atto 594+, yellow particles) represents efficient labelling of virions. Scale bar 5 μm. (B) Bar graph showing the number of mature and immature virions in viral samples prepared in absence (-) or presence of saquinavir (SQV) treatment. (C) Single particle tracking of mature (upper panel) and immature virions (bottom panel) upon saponin-induced membrane permeabilization. The micrograph on the left shows a double labelled HIV-1 mature particle (GFP+ Atto 594+) releasing the GFP content at ~240 s after saponin addition, as observed by a drop in green fluorescence intensity (right chart, upper panel). Membrane permeabilization in immature HIV-1 particles instead allows access to uncleaved Gag-GFP by NbA594, judged by an increase in red fluorescent intensity at ~200 s after saponin addition (right chart, bottom panel). Scale bar 5 μm. A.U.: arbitrary units.

**Figure S2.**
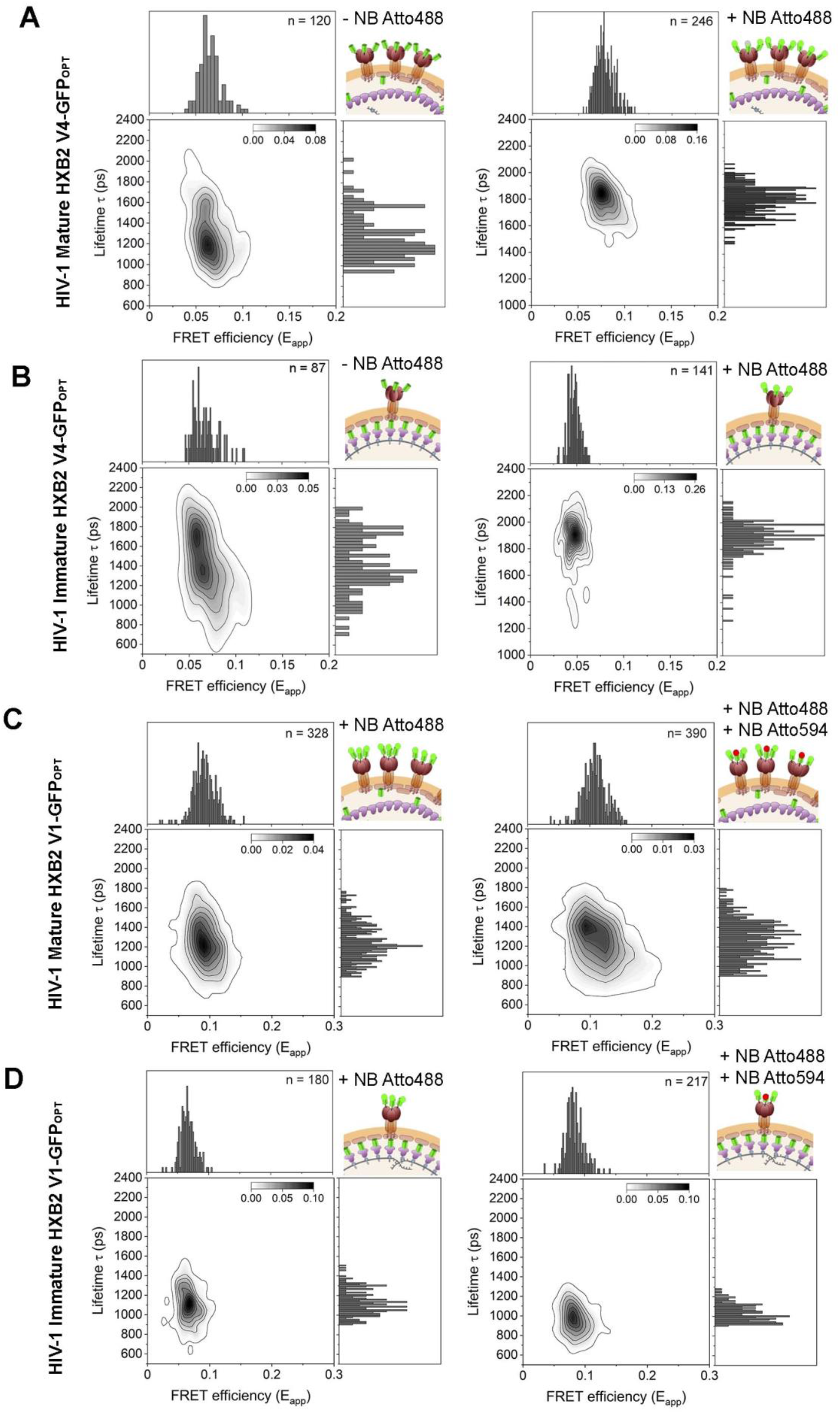
FRET negative controls for intramolecular and intermolecular HIV-1 Env conformations. Two-dimensional (2D) kernel probability graphs showing FRET (FRET efficiency, E_app_) vs FLIM (Lifetime, in ps) data. (A-B) HXB2 V4-GFP_OPT_ bearing Gag-GFP labelled without (left charts) or with (right charts) NbA488. Addition of the donor fluorophore, NbA488, induces a shift towards longer lifetimes, allowing both, to identify double labelled (A) mature and (B) immature particles (GFP+ NbA488+), and to define the negative signal for intramolecular FRET interactions (C-D) HXB2 V1-GFP_OPT_ virions bearing Gag-GFP labelled with the donor dipole, NbA488, in absence (left charts) or in presence of the acceptor dipole, NbA594 (right charts). Addition of NbA594, induces a shift towards higher FRET efficiencies in both, (C) mature and (D) immature particles below 0.23, which determines the threshold for intermolecular interactions observed in gp120 V4-labelled virions.

**Figure S3.**
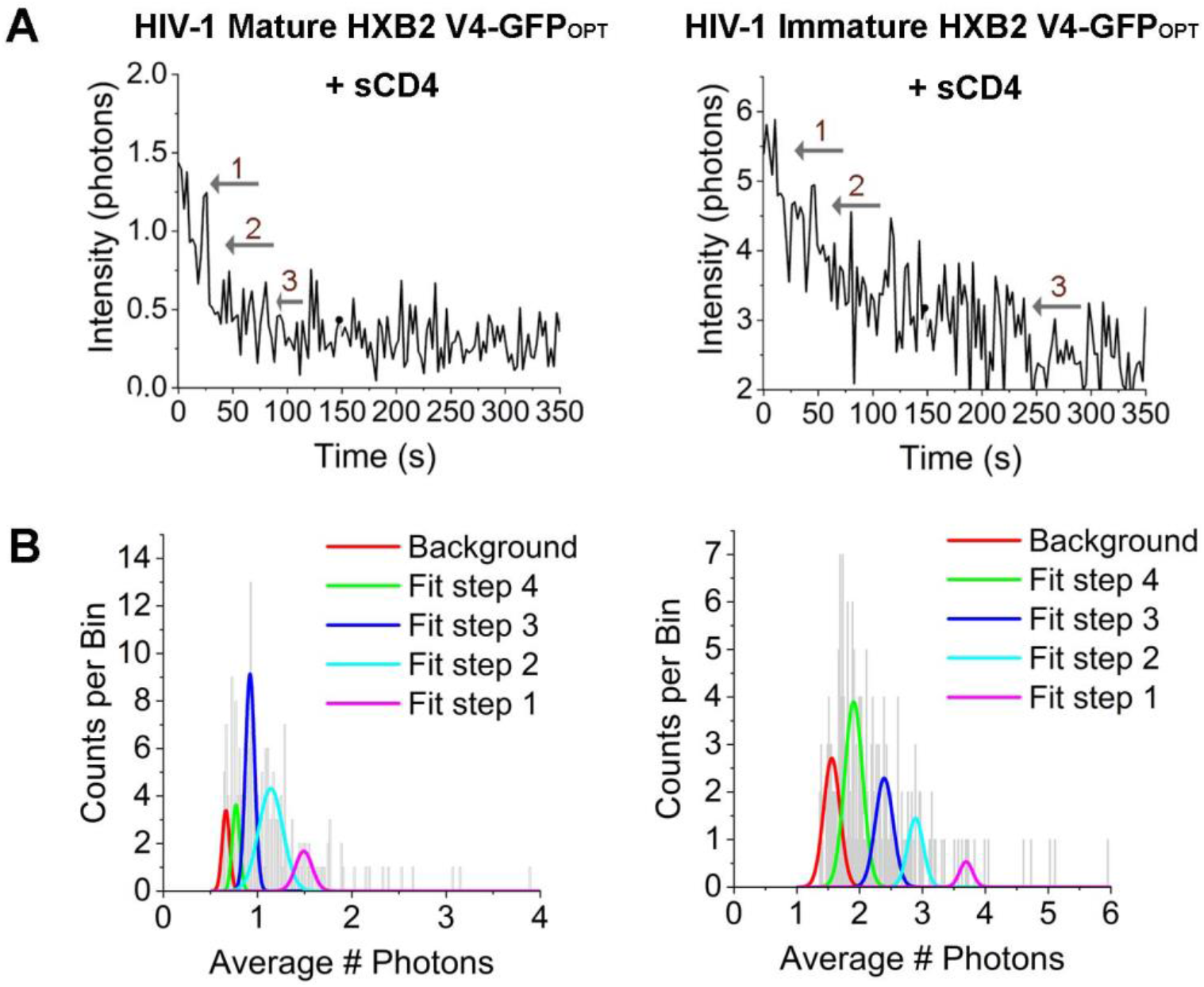
Single-molecule step photobleaching of HIV-1 Env in mature vs immature HXB2 V4-GFP_OPT_ pseudotyped virions in presence of sCD4. (A) Representative intensity traces for HIV-1 mature (left) and immature (right) HXB2 V4-GFP_OPT_ particles. Arrows point to single photobleaching steps detected. (B) Population histograms calculated from intensity traces were fitted into a multi-Gaussian distribution model to estimate the number of photobleaching steps for mature (left chart) and immature (right chart) viral particles. Χ^2^ values report the goodness of the fit.

**Figure S4.**
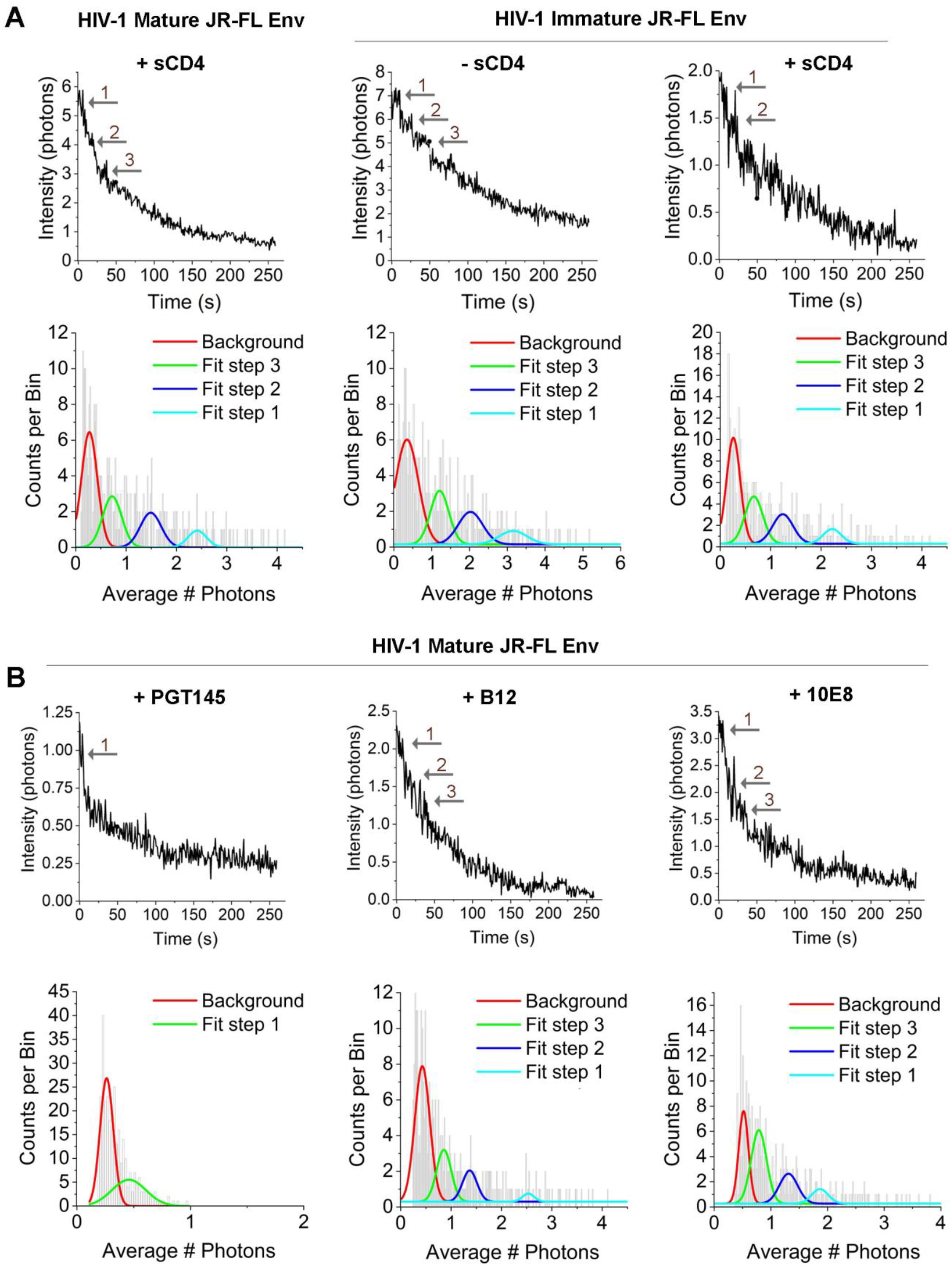
Single-molecule step photobleaching of HIV-1 JR-FL Env in mature vs immature virions in presence of sCD4 or neutralizing antibodies. (A) Representative intensity traces for HIV-1 mature (left) and immature (middle and right) HIV-1 JR-FL particles and the corresponding population histograms calculated from intensity traces. Histograms were fitted into a multi-Gaussian distribution model to estimate the number of photobleaching steps for mature (left chart) and immature (middle and right charts) viral particles in presence or absence of sCD4. Χ^2^ values report the goodness of the fit. Arrows point to single photobleaching steps. (B) Analysis as in (A) of HIV-1 mature JR-FL Env virions in presence of neutralizing antibodies PGT145, b12 and 10E8, targeting different regions of the Env glycoprotein.

**Figure S5.**
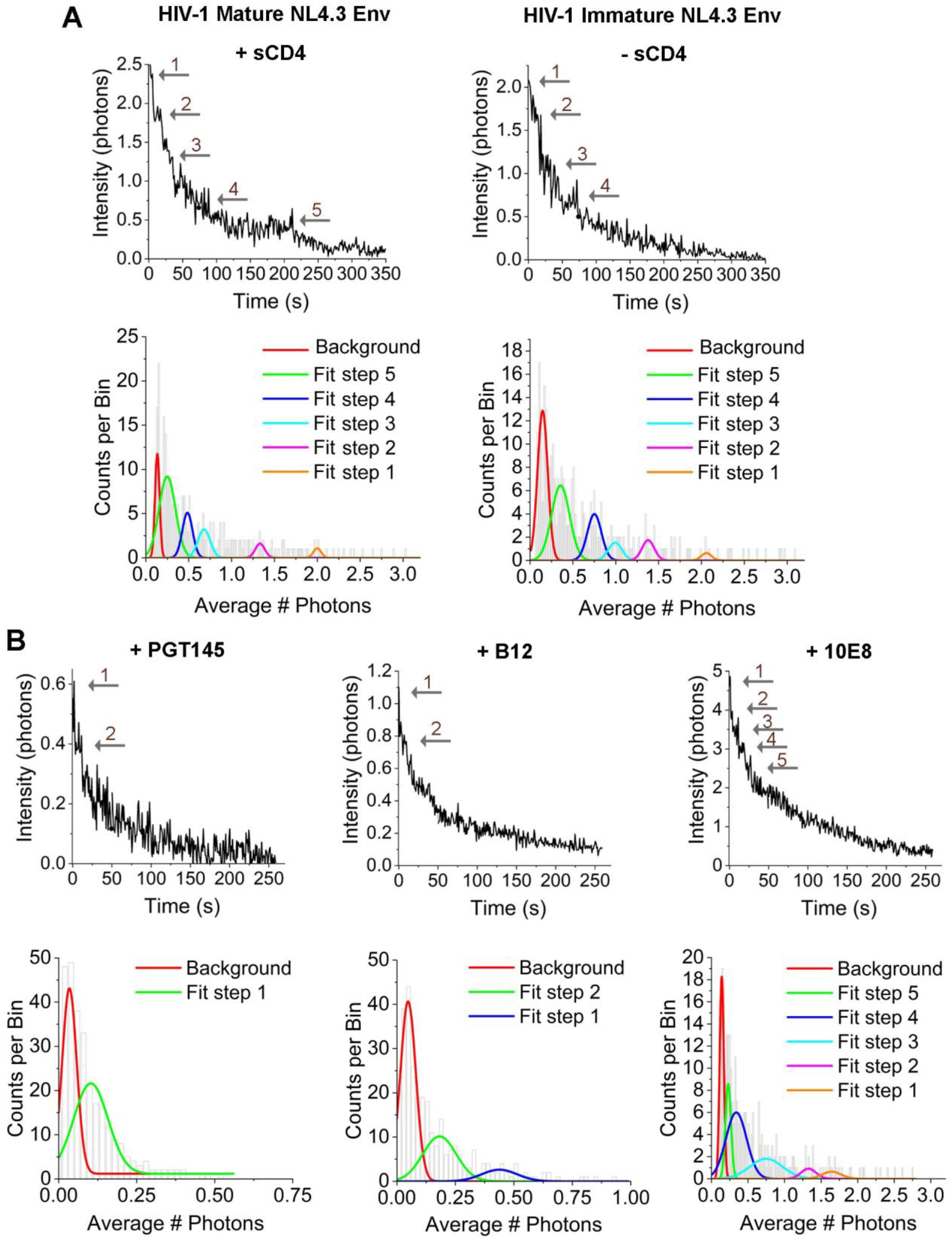
Single-molecule step photobleaching of HIV-1 NL4-3 Env in mature vs immature virions in presence of sCD4 or neutralizing antibodies. (A) Representative intensity traces for HIV-1 mature in presence of sCD4 (left) and immature in absence of sCD4 (right) HIV-1 NL4-3 particles and the corresponding population histograms calculated from intensity traces. Histograms were fitted into a multi-Gaussian distribution model to estimate the number of photobleaching steps for mature (left chart) and immature (right chart) viral particles. Χ^2^ values report the goodness of the fit. Arrows point to single photobleaching steps. (B) Analysis as in (A) of HIV-1 mature NL4-3 Env virions in presence of neutralizing antibodies PGT145, b12 and 10E8, targeting different regions of the Env glycoprotein.

